# Maximizing memory capacity in heterogeneous networks

**DOI:** 10.1101/2024.09.25.615056

**Authors:** Kaining Zhang, Gaia Tavoni

## Abstract

A central problem in neuroscience is identifying the features of neural networks that determine their memory capacity and assessing whether these features are optimized in the brain. In this study, we estimate the capacity of a general class of network models. Our derivation extends previous theoretical results, which assumed homogeneous connections and coding levels (i.e., activation rates of the neurons in memory patterns), to models with arbitrary architectures (varying constraints on the arrangement of connections between cells) and heterogeneous coding levels. Using our analytical results, we estimate the memory capacity of two types of brain-inspired networks: a general class of heterogeneous networks and a two-layer model simulating the CA3-Dentate Gyrus circuit in the hippocampus, known to be crucial for memory encoding. In the first case, we demonstrate that to maximize memory capacity, the number of inward connections and the coding levels of neurons must be correlated, presenting a normative prediction that is amenable to experimental testing. In the second case, we show that memory capacity is maximized when the connectivity and coding levels are consistent with the formation of memory “indices” in the Dentate Gyrus, which bind features in the CA3 layer. This suggests specific neural substrates for the hippocampal index theory of memory encoding and retrieval.

## I. INTRODUCTION

In a seminal paper [1], Hopfield introduced a type of artificial neural networks - inspired by models from statistical physics [2–4] - that can store memories and retrieve them from partial cues, providing a well-defined theoretical framework for the study of memory function in the brain [5–12]. The classic Hopfield model is a network of binary (active or silent) neurons that are potentially fully connected. Memories are stored by adapting the synaptic weights according to biologically inspired Hebbian plasticity rules [1, 13]. After this learning process, the memories are encoded in the matrix of synaptic connections and function as attractors or fixed points of the network dynamics: starting from a random activity configuration, the neurons with strong positive inputs become active while the other ones become silent, until the network converges to one of the memory patterns [5]. Memories can only be stored up to some maximum capacity. The quantification of the capacity of Hopfield networks has been the subject of an extensive literature. Early theoretical research demonstrated that the maximum number of random patterns that can be stored by a simple Hebbian learning rule is *C* ∼ 0.14 *N* [1, 14, 15]: the capacity increases linearly with the number of neurons. Gardner subsequently introduced a method to compute the maximal memory capacity, independent of the learning rule used [16–18]. This upper bound is *C* = 2*N* for uncorrelated random patterns and is attainable with a local learning algorithm, also introduced in [16]. This research focused on abstract networks of fully connected binary neurons with the same coding levels, i.e., the same activation probabilities in the memory patterns. Other studies have considered diluted Hopfield networks, in which connections are randomly dropped with a fixed probability [6, 19–21], and fully or sparsely connected homogeneous Hopfield networks with discrete synapses [22]. A few studies have addressed the conditions for memory retrieval in heterogeneous networks [6, 23, 24], but these extensions are limited to symmetric graphs (where neurons are reciprocally connected) and symmetric Hebbian learning. All these analyses also maintained homogeneous coding levels across all neurons.

Through this fundamental theoretical research, powerful tools have been developed to study memory in neural networks. However, the application of these tools to biologically realistic cases has remained a relatively under-explored field. Of note, biological networks are neither fully nor homogeneously or symmetrically connected and neurons may have different coding levels. In the spirit of gaining insights into memory function in real brains, we derive an analytical solution for the maximal memory capacity of a general class of networks with arbitrary architectures, i.e., any possible arrangements of connections between cells, and arbitrary distributions of coding levels across neurons. Our analysis is crucial for understanding the influence of various structural and neurophysiological constraints on the ability of networks to store information.

The problem of estimating the memory capacity of Hopfield networks is tightly linked to the perceptron capacity problem. A perceptron is a simple feedforward network with a binary output unit that linearly separates N-dimensional input patterns into predefined classes. The perceptron capacity is the maximum number of input patterns that can be linearly assigned to their classes or labels. The methods developed to analyze perceptrons and their capacity can often be employed to study Hopfield networks, and vice versa [16, 25–31]. Previous research has analyzed perceptrons and Hopfield networks with uncorrelated and correlated units or patterns [16, 27, 32, 33], correlated input-label pairs [34], and other relevant classification and learning scenarios beyond storage capacity [26, 32, 33]. However, a formula for the perceptron capacity in the case of independent but otherwise arbitrarily distributed input states is still missing. We begin our study by deriving this formula and demonstrate that it generalizes to a broad class of perceptrons with correlated input states. Using this result, we then calculate the memory capacity of heterogeneous Hopfield networks with arbitrary architectures and neuron coding levels. We find that the capacity of perceptrons is unaffected by the distribution of input states and depends only on the mean value of the labels (or coding level of the output unit). Extending the calculation to networks, we find that the capacity is a function of the coding level and number of inward connections of each cell.

Finally, we use these analytical results to study the memory capacity of biologically inspired networks. We first consider a general class of heterogeneous networks in which the number of inward connections and coding levels of the neurons are not constant but are randomly sampled from distributions with different variances. These variances determine the degrees of heterogeneity in the networks. We find that increasing heterogeneity generally reduces the network memory capacity, except when coding levels and numbers of inward connections are correlated: if neurons with more inward connections have proportionally higher coding levels, the memory capacity of the network is maximized. Secondly, we consider a network composed of two reciprocally connected, but internally disconnected, layers. We assume that one layer (the “feature or visible layer”) stores the memory features, while the other (the “memory or hidden layer”) forms an internal representation of the memory patterns [6, 8, 35–37]. We use this network architecture to represent, at a high level, the CA3-DG (Cornu Ammonis 3 - Dentate Gyrus) circuit in the hippocampus, during the initial memory encoding stage. We show that the capacity for storing information is maximized in this model when the hidden (DG) units have sparse coding levels and the learned inter-layer connectivity is consistent with the formation of memory “indices”, i.e., neural units in the hidden layer that selectively receive inputs from and point to the features of specific memories in the visible layer (CA3). This network configuration is believed to support the pattern separation and pattern completion functions of the hippocampus, which are important for memory retrieval [38–42].

## II. RESULTS

### A. Capacity of perceptrons with arbitrarily-distributed input states

A perceptron (Fig. 1A) is a simple feedforward network with one output unit that responds to *N* input units according to the rule:

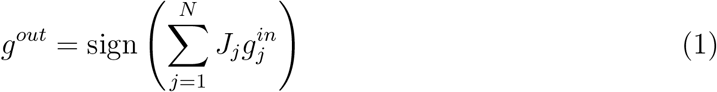

where *g*^*out*^ and 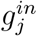 represent the output and the *j*-th input state, respectively, and {*J*_*j*_} are the synaptic weights: the output state is the sign of the total input to the output unit.

**FIG. 1.**
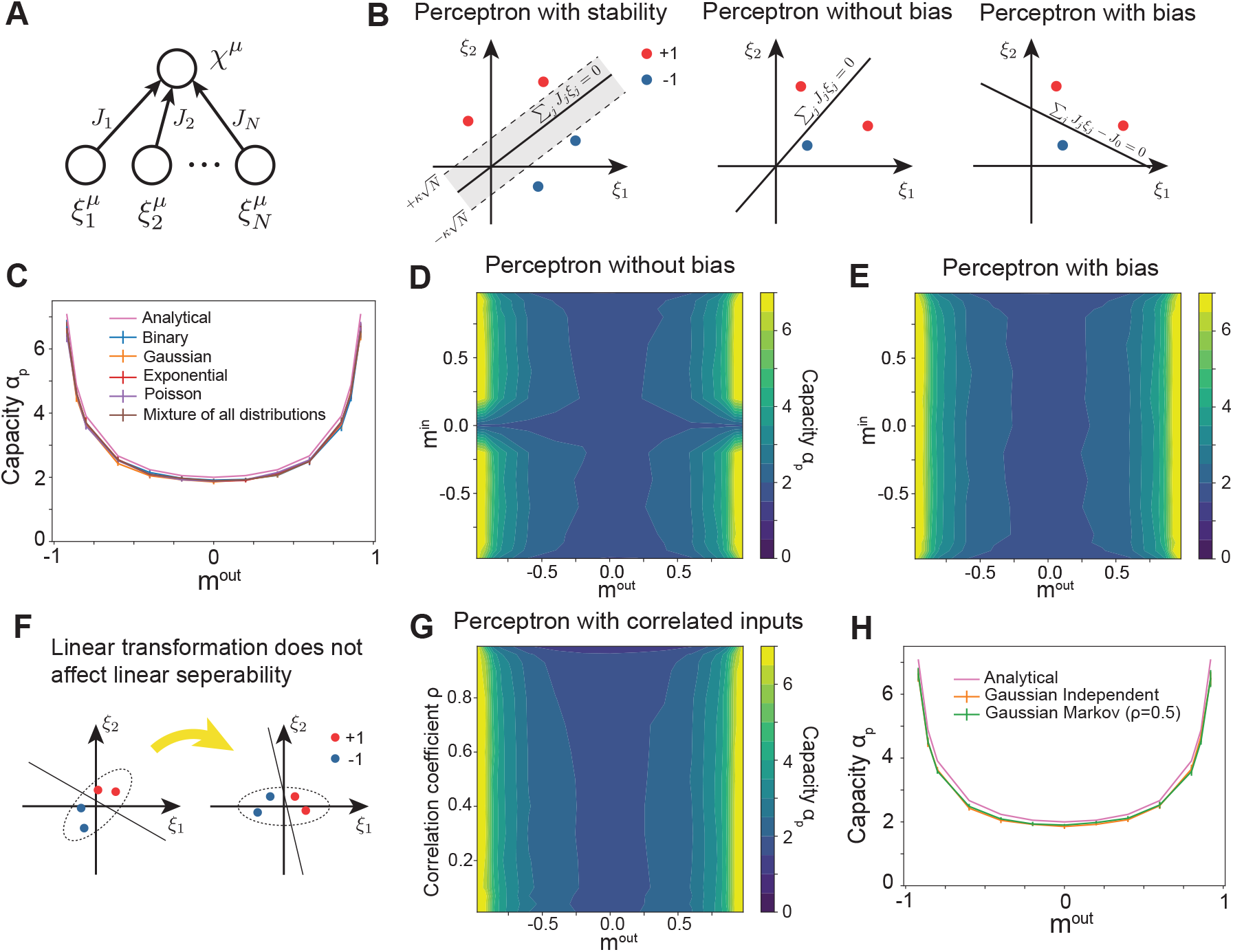
Perceptron capacity. **A**. A perceptron is a feedforward network that stores pattern-label pairs in the weights {*J*_*j*_}; inputs 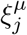 can have arbitrary (independent) distributions with distinct finite means and variances and labels *χ*^*µ*^ are binary. **B**. A perceptron is a linear classifier. Axes: input states; dots: input patterns (here *N* = 2 for simplicity). Within capacity, the perceptron can find a hyperplane (solid line) that separates the patterns according to their labels (red and blue). Left: to enforce classification stability against noise, the patterns must be at a minimum distance (set by *κ*) from the hyperplane (dashed lines). Middle: for perceptrons without bias, the decision hyperplane must pass through the origin. Right: for perceptrons with bias, the decision hyperplane is not constrained to pass through the origin, improving linear separability in low dimension. **C**. Maximal capacity of a perceptron as a function of the output mean *m*^*out*^; analytical solution, Eqs. (3) (magenta), and mean ± std over 10 numerical estimates (N=1000, Methods) for perceptrons with bias and different input distributions (different colors). **D**. Numerical estimates of the maximal capacity of a perceptron without bias. Input (output) states are independently sampled from a binary distribution with mean *m*^*in*^ (*m*^*out*^). **E**. Same as (D) for a perceptron with bias. In the large *N* limit, the capacity depends only on *m*^*out*^ (not on *m*^*in*^). Slight variations in capacity with *m*^*in*^ are due to finite-size effects. **F**. Linear separability of the input patterns is not affected by any linear transformation of the patterns. **G, H**. The maximal capacity is not affected by correlations in the inputs (as long as these correlations can be removed by a linear transformation of the patterns in the input space). **G**. Input states are sampled from a Gaussian Markov chain (Methods) and numerical estimates of the maximal capacity are shown as a function of *m*^*out*^ and the correlation coefficient *ρ* between adjacent input states 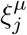 and 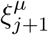. **H**. Analytical solution as a function of *m*^*out*^, Eqs. (3) (magenta), coincides with numerical estimates for uncorrelated (*ρ* = 0, orange) and correlated (*ρ* = 0.5, green) input states.

A perceptron can learn to associate N-dimensional input patterns 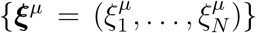with specified binary classes or labels {*χ*^*µ*^ = ±1}. The input-label pairs are learned by adjusting the synaptic weights {*J*_*j*_} so that they satisfy the classification constraint

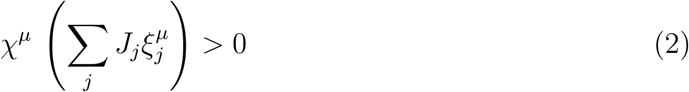

for every input pattern indexed by *µ*. Constraints (2) define a hyperplane in the input space {*ξ*_*j*_} (*j* = 1, …, *N*) that separates the patterns labeled as +1 from those labeled as −1 (Fig. 1B, left, solid line). A more restrictive condition can be enforced, requiring that a minimum distance exists between the patterns and the hyperplane separating the classes. This condition ensures that the classification is stable against a proportional amount of noise. The minimum distance between the patterns and the hyperplane is determined by a stability parameter *κ >* 0 (Fig. 1B, left, dashed lines).

The capacity *C*_*p*_ is the maximum number of input patterns that the perceptron can learn to classify with probability 1, according to the conditions described above. We compute the normalized capacity, *α*_*p*_ = *C*_*p*_*/N*, in the general case of input-pattern states 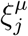 that are *independent* but otherwise *arbitrarily distributed*, each with a distinct and *finite mean m*_*j*_ and *finite variance* 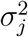. Different means correspond to different activation probabilities of the units in the input patterns, i.e., different neuron coding levels. We assume that the labels follow a Bernoulli distribution with *p*(*χ*^*µ*^ = 1) = (1 + *m*^*out*^)*/*2 and *p*(*χ*^*µ*^ = −1) = (1 − *m*^*out*^)*/*2; *m*^*out*^ is the output (or label) mean, which can be different from the input means {*m*_*j*_}. We thus address the memory capacity problem in a more general class of perceptrons than in [16], where the maximal capacity was obtained for *independent, binary*, and *identically distributed* input-pattern states 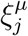 and output labels *χ*^*µ*^.

To calculate the capacity of this general class of perceptrons, we use a method pioneered by Gardner, based on estimating (for large *N*) the typical volume of the synaptic weights that satisfy the classification constraints for all the input patterns [16, 17, 43] (details in Appendix A). The capacity is maximized when this volume shrinks to zero, i.e., no input pattern can be added without breaking the constraints.

In the *κ* = 0 case, the solution to the classification constraints (2) is completely determined by the following joint equations, which can be evaluated numerically (Appendix A and Methods):

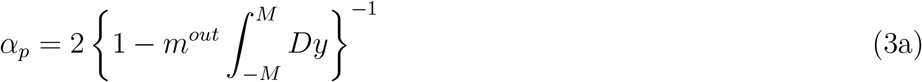

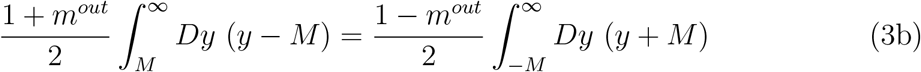

The integration limits (±*M*) in Eq. (3a) are given by Eq. (3b) when *m*_*j*_ ≠ 0 for at least one input unit. When *m*_*j*_ = 0 for all input units, *M* = 0 and *α*_*p*_ = 2.

The parameters 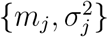, or any other details of the input-state distributions, do not factor into the solution, Eqs. (3). Thus, the perceptron capacity is independent of the statistics of input-pattern states and depends only on the output mean activation *m*^*out*^ (the average value of the labels). The capacity is maximized at extreme (low or high) values of *m*^*out*^ (Fig. 1C, magenta curve). The only exception is when the distribution of input states is centered at the origin (*m*_*j*_ = 0 for all *j*): as mentioned above, at this singular point, the maximal capacity is *α*_*p*_ = 2, regardless of *m*^*out*^ (Fig. 1D).

Enforcing classification stability against noise (*κ >* 0) does not change these results, except for an overall reduction in capacity as *κ* increases (Supplemental Fig. 1).

#### Perceptrons with bias and correlated input units

A more general definition of the perceptron includes a bias term corresponding to an activation threshold of the output unit:

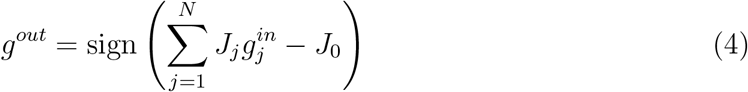

Introducing the bias term *J*_0_ is equivalent to adding a formal input unit with weight *J*_0_ and constant state 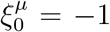 for every pattern *µ* [16]. This is equivalent to sampling an additional state from a binary distribution with 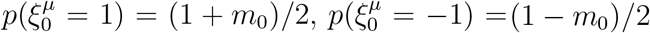, where *m*_0_ → −1.

Since the perceptron capacity does not depend on the distributions of input states, and the increase in dimension from *N* to *N* + 1 is irrelevant in the large *N* limit, the capacity formula (3) is still valid in this case. Due to the additional formal unit with *m*_0_ = −1, the same formula also applies when *m*_*j*_ = 0 for *j* = 1, …, *N* in perceptrons with bias, removing the singularity present in the case without bias. In summary, in the large *N* limit, the capacity of the perceptron with bias is determined solely by the mean value of the labels *m*^*out*^ and is independent of the input statistics. We confirm this result in numerical estimates of the solution (Fig. 1CE).

The general equivalence between the capacities of perceptrons with and without bias breaks down in low dimensions (small *N*). This is illustrated geometrically in Fig. 1B for *N* = 2. For perceptrons without bias, the hyperplane (Σ _*j*_ *J*_*j*_*ξ*_*j*_ = 0) separating the input patterns into binary classes must intersect the origin of the input space (Fig. 1B, left and middle); on the contrary, the bias term allows the hyperplane (Σ_*j*_ *J*_*j*_*ξ*_*j*_ = *J*_0_) to be shifted and tilted in such a way that it can better separate the input patterns without the restriction of passing through the origin (Fig. 1B, right). While in high dimension adding a single constraint - the intersection of the hyperplane with the origin - has a vanishing impact on classification performance (except when all *m*_*j*_ = 0), in low dimension it can substantially decrease the perceptron capacity.

Our solution for the perceptron capacity further generalizes to the case of *correlated input states*, even if Eqs. (3) are derived under the assumption that the input states 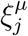 are independent. This invariance property follows from the fact that linear separability is not affected by any linear transformation of the input patterns. Thus, the capacity of the perceptron (with bias) remains unchanged when the input units are correlated, as long as these correlations can be removed by a linear transformation of the patterns in the input space (Fig. 1F). We validate this argument by numerically estimating the capacity of a perceptron with input states sampled from a Gaussian Markov chain, so that each unit is correlated with its neighbors (similar to the case considered in [27], details in Methods). The estimated capacity aligns with our analytical and numerical solutions for uncorrelated inputs (Fig. 1GH).

### B. Capacity of Hopfield networks with arbitrary architectures and neuron coding levels

We can now use the perceptron results to obtain the capacity of Hopfield neworks with heterogeneous architectures (arbitrary arrangements of connections between cells) and coding levels (different activation probabilities of the neurons in the memory patterns), Fig. 2A. We define a network architecture - or structure - in terms of a binary matrix *J*^*S*^ with 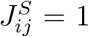 if neuron *j* is connected to neuron *i* (in the *j* → *i* direction) and 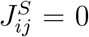 if the neurons are not connected in that direction. The matrix can be asymmetric and we assume that neurons do not connect to themselves 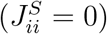. We further define *P* patterns 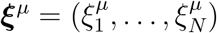 to memorize, such that 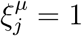 (active) with probability *p*_*j*_ and 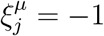 (inactive) with probability 1 − *p*_*j*_. Thus, *p*_*j*_ represents the coding level of neuron *j* or the expected fraction of memory patterns in which that neuron is active; unlike previous studies, our definition allows for different coding levels for different neurons (Fig. 2A) [7, 22, 23, 27, 44].

**FIG. 2.**
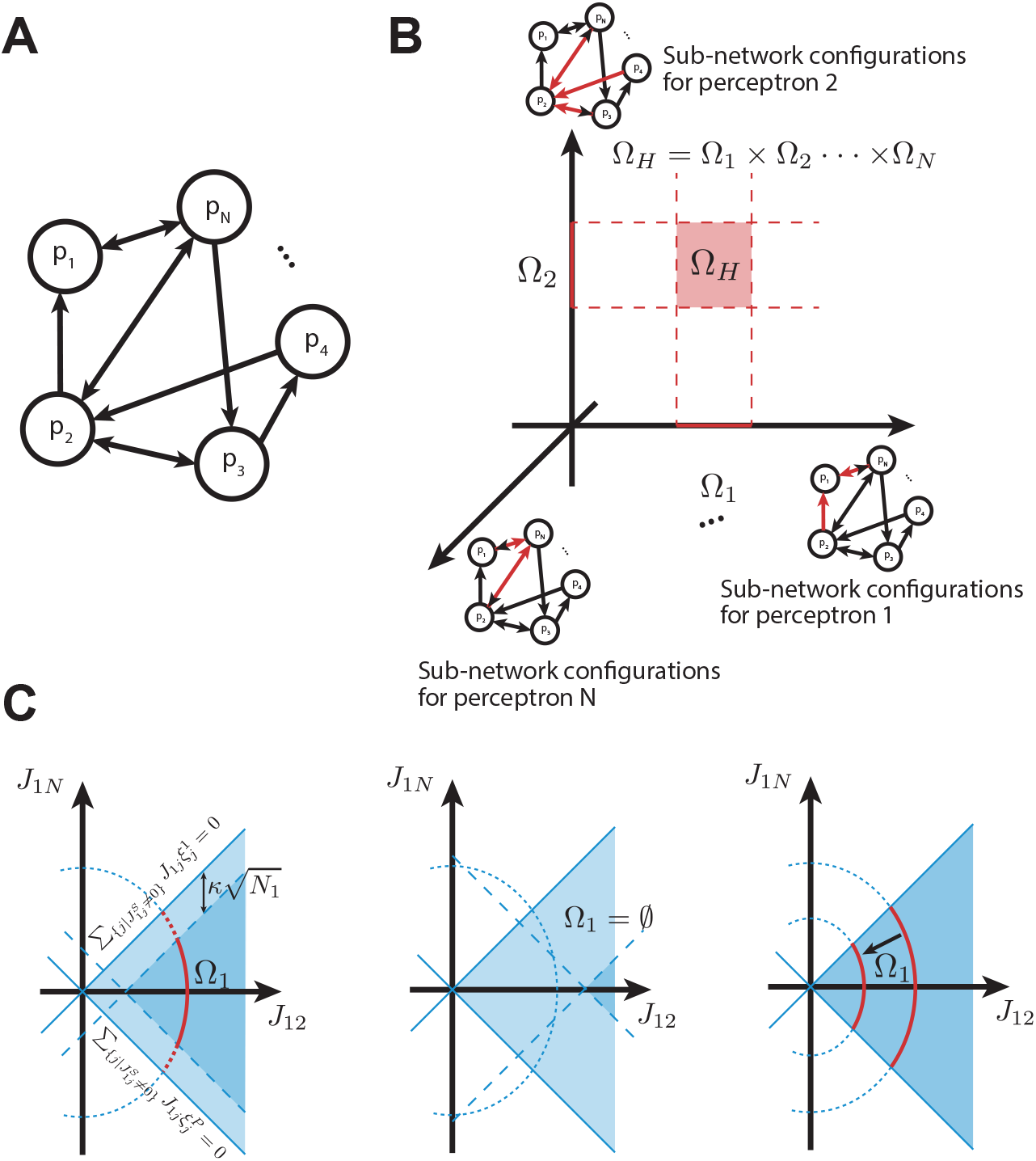
Calculating the capacity of heterogeneous Hopfield networks. **A**. A heterogeneous Hopfield model is defined by an arbitrary architecture, or arrangement of directed connections 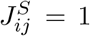 (arrows), and arbitrary neuron coding levels {*p*_*i*_}. **B**. Sketch of the so-lution set Ω_*H*_ for the Hopfield model, i.e., the ensemble of network configurations {*J*_*ij*_} that satisfy the fixed-point conditions (5) for the *P* memory patterns. Ω_*H*_ is the product of the solution sets Ω_*i*_ for the *N* independent perceptrons defined on the network. Ω_*i*_ is the ensemble of sub-network configurations for perceptron *i* (red arrows) that satisfy the subset of conditions (5) for that perceptron. The product Ω_1_ × · · · × Ω_*N*_ is defined by all possible combinations of solutions (sub-network configurations) in the different sets. Red square: projection of Ω_*H*_ onto the two-dimensional plane (Ω_1_, Ω_2_). **C**. A close examination of the solution set Ω_1_ for the sub-network corresponding to the 1st perceptron; the sketch is drawn for *N* = 3. Left: Ω_1_ con-sists of sub-network configurations 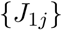 (with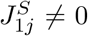) that satisfy the fixed-point conditions (5) for *i* = 1; *µ* = 1, …, *P* and the elliptic weight regularization 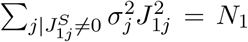. Each memory pattern constrains the solution space to be on one side of a hyperplane (a half-space); *P* memories define *P* hyperplanes (solid blue lines) and corresponding intersection of half-spaces (light blue region). The weight regularization constrains the solution space to lie on the ellipse (dotted blue line). Altogether, these conditions constrain the solution space (Ω_1_) to the red arc. Enforcing stability of the memories against noise corresponds to shifting each hyperplane by 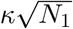, thus shrinking the intersection of half spaces (dark blue region) and the solution set (solid red line). Middle: Increasing *κ* can make the intersection of half-spaces (dark blue area) fall outside the elliptic region: the solution set Ω_1_ vanishes and the memory capacity decreases. Right: When *κ* = 0, the elliptic constraint contributes to defining Ω_*H*_ (different elliptic regularizations yield different solution sets, red arcs) but cannot make the solution set vanish. Thus, the maximal capacity is unaltered by the weight regularization when *κ* = 0.

The same activation rule used for the perceptron output, Eq. (1), defines a dynamics on Hopfield networks, whereby any given neuron is activated if it receives a positive total input from the other cells and it is inactivated if it receives a negative total input. There are fixed points in this dynamics: at those points, the state of every neuron has the same sign as its total input. The fixed points are memories stored in the network, which can be retrieved from partial cues - representing corrupted patterns - through the same dynamics. Thus, to store *P* memory patterns 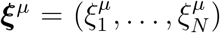, the network weights {*J*_*ij*_} need to satisfy the following fixed-point conditions:

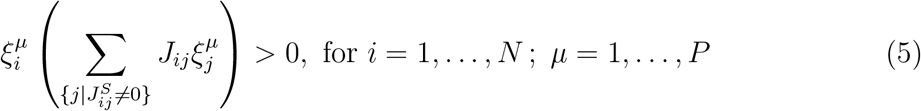

It is straightforward to see that the problem of storing memories in a Hopfield network with *N* neurons factorizes into *N* independent perceptron classification problems (Fig. 2B): each neuron *i* in the Hopfield network can be thought of as the output unit of one perceptron, with label 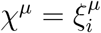, and all the neurons that have connections to it can be thought of as the input units for that perceptron, with pattern states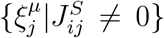; repeating this mapping for every neuron defines *N* perceptrons. The inequalities (5) are precisely the classification constraints (2) applied to all *N* perceptrons. Thus, the solution set for the Hopfield model, that is, the set of network configurations {*J*_*ij*_} with structure *J*^*S*^ that meet all the inequalities (5), is given by all possible combinations of *N* independent sub-network configurations that meet the classification constraints for the different perceptrons. In mathematical language, the solution set Ω_*H*_ for the Hopfield model is the Cartesian product of the solution sets Ω_*i*_ for the *N* perceptrons (Fig. 2B):

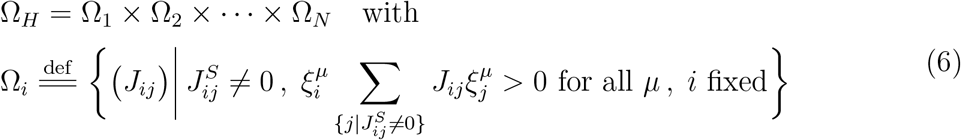

We define the memory capacity of the Hopfield network, *C*_*H*_ = *α*_*H*_ *N*, as the maximum number of patterns for which *all* the *N* perceptrons satisfy the fixed-point conditions with probability 1. Above this extreme number, there exists a set of realizations of the patterns with non-zero measure that violate the constraints in at least one perceptron (denoted by *i*). For those realizations of the patterns, the volume of the solution set Ω_*i*_, and thus the volume of Ω_*H*_, shrinks to zero (Appendix A). It follows that the memory capacity of the network, as we defined it, coincides with that of the perceptron with the smallest maximal capacity. Thus, for a heterogeneous Hopfield model, we obtain:

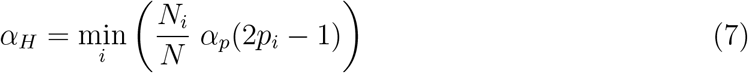

where 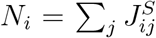 is the number of inward connections (the in-degree) of neuron *i* and *α*_*p*_(2*p*_*i*_ − 1) = *C*_*p*_*/N*_*i*_ is the normalized capacity of the perceptron with output neuron *i*.

The latter is a function of *m*_*i*_ = 2*p*_*i*_ − 1, the expected state of neuron *i* in the memory patterns (replacing *m*^*out*^ in Eqs. (3)).

This formula generalizes classical results on the capacity of Hopfield models to heterogeneous networks, accommodating arbitrary constraints on the architecture as defined by *J*^*S*^, and varying neuron coding levels as defined by the different *p*_*i*_.

Similarly to the perceptron, a more restrictive condition can be enforced, requiring the memory patterns to have sufficiently wide basins of attraction. This condition ensures that the memories can be retrieved starting from a relatively wide neighborhood of initial configurations, making them stable against a proportional amount of noise. The width of the basins of attraction of the memories is controlled by a stability parameter *κ* [7, 16]. Setting *κ >* 0 is equivalent to enforcing classification stability in each of the *N* perceptrons defined on the Hopfield network (Fig. 2C, left). Thus, the network capacity for *κ* ≥ 0 is given by Eq. (7) with *α*_*p*_ replaced by the capacity of the perceptrons with stability *κ* (derived in Appendix A). Increasing *κ* improves memory retrievability while reducing the memory capacity of the network (Fig. 2C, middle).

When *k* = 0, the capacity result applies to any Hopfield network as defined above; on the contrary, under the stronger condition *κ >* 0, the result holds true only for a class of Hopfield networks that satisfy the weight regularization 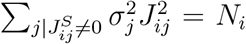 for every neuron *i*, where 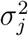 denotes the variance of the states of unit *j*. This regularization, which can be thought of as a normalization of the postsynaptic input ranges of the neurons, is utilized in the capacity calculation for any *κ* ≥ 0 but does not impact the result when *κ* = 0 (Fig. 2C, right, and Appendices A, B).

### C. Biologically relevant case studies

In the spirit of gaining insights into the memory capacity of real neural networks, we apply our analytical results to two biologically relevant case studies: a general class of heterogeneous networks and a class of two-layer memory networks.

#### Correlation between neuron in-degrees and coding levels maximizes the memory capacity of heterogeneous networks

Neurons in the brain can have different activation probabilities within memory patterns (coding levels), different numbers of inward connections (in-degrees), and different numbers of outward connections (out-degrees). Thus, memory networks in the brain are largely heterogeneous.

Our analytical formula (7) demonstrates that the maximum number of patterns that can be stored as fixed points in heterogeneous networks depends only on the neuron coding levels *p*_*i*_ and in-degrees *N*_*i*_; conversely, the neuron out-degrees do not influence memory capacity. Here, we assess how the joint statistic of the two relevant parameters affects capacity and under which conditions such capacity is maximized.

We generate networks with different distributions of coding levels and in-degrees: specifically, we vary the standard deviation of coding levels and inward-connection probabilities across neurons while keeping their average values constant (Methods). Increasing the standard deviation of either parameter increases heterogeneity (Fig. 3A). In heterogeneous networks, the in-degrees and coding levels of the neurons can be independent, positively correlated or negatively correlated (Fig. 3B-D). We analyse below the effects of heterogeneity in these three different cases.

**FIG. 3.**
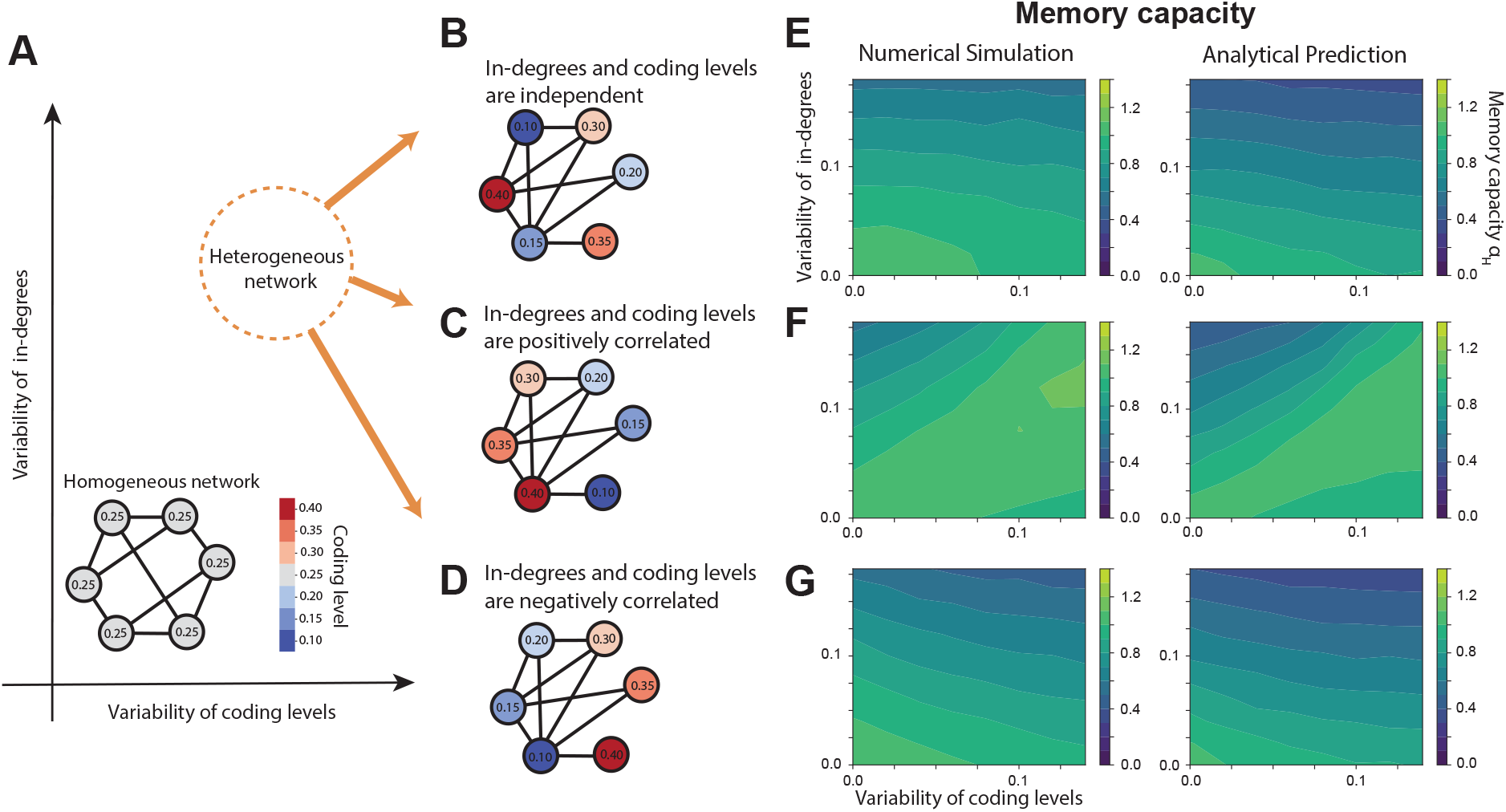
Memory capacity is strongly influenced by the joint statistic of neuron indegrees and coding levels. **A**. We generate networks with different levels of heterogeneity by changing the variability of coding levels and in-degrees across neurons (specifically, the standard deviations of coding levels and inward-connection probabilities, respectively); the means over neurons of both parameters are held constant across networks (Methods). Networks with low variability of both coding levels and in-degrees are homogeneous, whereas networks with high variability in these parameters are heterogeneous. **B-D**. In heterogeneous networks, neuron in-degrees and coding levels can be independent (**B**), positively correlated (**C**) or negatively correlated (**D**). **E-G**. Memory capacity as a function of the variability of in-degrees and coding levels, in the biologically plausible (*p*_*i*_ *<* 0.5) regime: numerical (left) and analytical (right) estimates. **E**: When in-degrees and coding levels are independent, memory capacity decreases as the variability of either parameter increases. **F**: When in-degrees and coding levels are positively correlated, the capacity of heterogeneous networks (with high variability of both parameters) is similar to the capacity of homogeneous networks (with low variability). **G**: When in-degrees and coding levels are negatively correlated, memory capacity undergoes the steepest decrease with increasing network heterogeneity.

When in-degrees and coding levels are independent, increasing the standard deviation of either parameter decreases the memory capacity of the network (Fig. 3E). This result can be understood as follows: the network capacity (normalized by the number of neurons, Eq. (7)) is the minimum of the weighted perceptron capacities 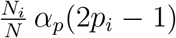; when increasing the variability of *N*_*i*_ or *p*_*i*_ across neurons, the *variance* of the weighted perceptron capacities increases, while the *mean* remains more stable. Consequently, the *minimum* of the weighted perceptron capacities becomes lower, which makes the network capacity decrease.

This capacity reduction in heterogeneous networks is mitigated when the distributions of in-degrees *N*_*i*_ and coding levels *p*_*i*_ are such that the weighted perceptron capacities are roughly constant across the neurons. In the regime of low coding levels, *p*_*i*_ *<* 0.5 (corresponding to *m*_*i*_ *<* 0), the perceptron capacity decreases with increasing *p*_*i*_ (Fig. 1C). Thus, for the weighted perceptron capacity to be roughly constant, neurons with higher *p*_*i*_ should have higher *N*_*i*_. In other words, neurons with higher coding levels should have denser inward connections. The effects of a positive correlation between *p*_*i*_ and *N*_*i*_ on the network memory capacity are shown in Fig. 3F. Compared to Fig. 3E, the capacity remains high in the heterogeneous region, when both in-degrees and coding levels are highly variable in the network.

This result is reversed when neurons with higher coding levels have sparser inward connections (negative-correlation case): as expected from the above argument, the network capacity decreases even faster in this condition than in the independent case, as the heterogeneity increases (Fig. 3G).

As a control, we confirmed that memory capacity is insensitive to neuron out-degrees, regardless of the correlation pattern between out-degrees and coding levels (Supplemental Fig. 2). Furthermore, enforcing memory robustness (*κ >* 0) does not qualitatively change our conclusions (Supplemental Fig. 3).

So far, we have focused on the regime of low coding levels (*p*_*i*_ *<* 0.5), which is more energy-efficient [45, 46] and biologically plausible, as experimental data show that neurons tend to exhibit sparser firing rates in response to memorized objects compared to novel ones [47–50]. In the opposite (less biologically plausible) regime, in which all *p*_*i*_ *>* 0.5, results are reversed: the network capacity is maximized when in-degrees and coding levels are negatively correlated, due to the non-monotonic trend of perceptron capacity with *p*_*i*_ (Fig. 1C). In this regime, the perceptron capacity increases with *p*_*i*_, requiring high coding levels to be associated with low in-degrees to maintain the weighted perceptron capacity roughly constant across neurons (Supplemental Fig. 4).

#### Formation of memory “indices” in a two-layer network optimizes capacity

We now consider a network model with two layers, representing two distinct brain areas necessary for memory function (Fig. 4A) [38, 41, 51–53]. The network structure, defined by *J*^*S*^, is such that reciprocal connections are present between the two layers but not within each layer. This type of bipartite architecture is closely inspired by the Bidirectional Associative Memory (BAM) model introduced by Kosko [35], and more loosely by Restricted Boltzmann Machines (RBM) and other modern Hopfield networks used to simulate memory systems [6, 8, 36, 37]). Compared to the BAM model, the synaptic connections do not need to be symmetric in our case. In our model, all units are subject to the same activation rule, Eq. (1), and the input patterns are stored across both layers. The two layers differ only in the number of neurons and their coding levels.

**FIG. 4.**
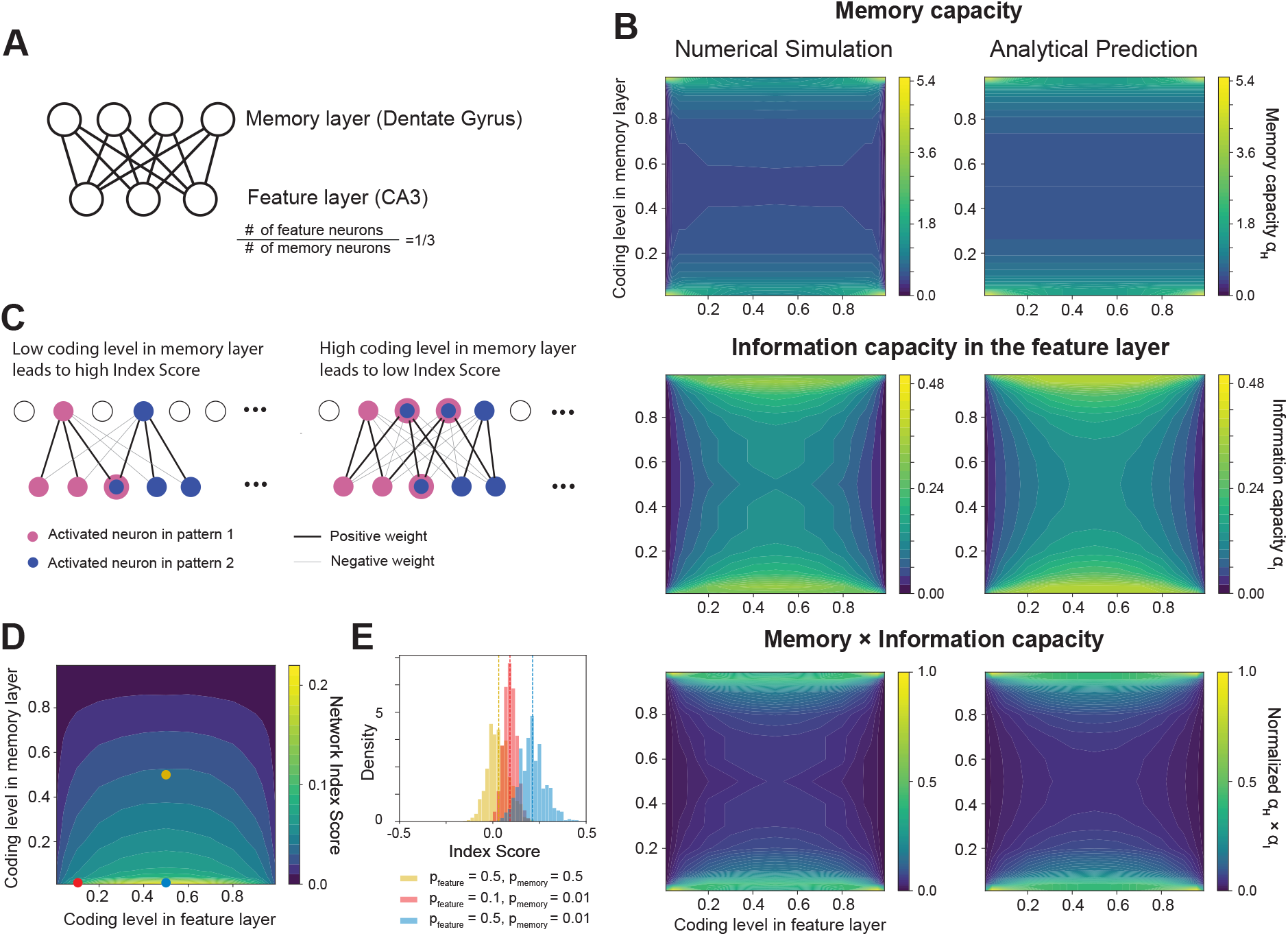
Capacity of two-layer networks and alignment with hippocampal index theory (HIT). **A**. Sketch of a two-layer network with reciprocal connections between the two layers and no connection within each layer; the visible or feature layer has fewer units than the hidden or memory layer. Here, we set the numbers of units to 150 and 450, respectively, with a 1:3 ratio to simulate the CA3-DG circuit in the hippocampus. **B**. Memory (top), information (middle), and memory × information (bottom) capacity (the latter normalized by the maximum value) for the two-layer network shown in (A), as a function of the coding levels in the feature and memory layers: numerical (left) and analytical (right) estimates. Both memory and information capacities are more sensitive to the coding level in the memory layer. When this is low or high, both capacities are maximized. **C**. The Index Score quantifies how well a memory unit active in a given memory pattern selectively binds the features of that pattern: examples of memory units with a high (left) and a low (right) Index Score, corresponding to conditions of low or high coding levels in the memory layer, respectively. **D**. The Network Index Score (average score of the active memory units in the stored patterns) varies with coding levels in the feature and memory layers. It is maximum when the coding level in the memory layer is low, which coincides with a condition of maximum memory and information capacity. **E**. Distributions of Index Scores for the active memory units in the stored patterns are shown for different coding levels: both intermediate in the memory and feature layers (yellow), low and intermediate, respectively (blue), or both low (red), corresponding to the same-colored dots in (D). When the coding level in the memory layer is low, the distributions of Index Scores are entirely above zero, indicating an indexing function for the memory units, as posited by HIT.

Inspired by theories of memory encoding and consolidation in the brain, we interpret one of the two partitions, the “visible or feature” layer, as the storage location for the informational content of memories [38, 41, 51–58]. We interpret the other partition, the “hidden or memory” layer, as a memory support system that can potentially bind the feature neurons and stabilize the patterns as fixed points of the dynamics. We use this model as a high-level description of a hippocampal network crucial for memory function: the Cornu Ammonis 3 - Dentate Gyrus (CA3-DG) circuit. Empirical evidence suggests that, within the hippocampus, CA3 may store the informational content of experiences (serving as the feature layer), while the DG may act as a teacher, providing the initial signal to bind the features in CA3 (serving as the memory layer) [38–40, 42]. Here, we focus on the initial stage of memory encoding. Previous computational and experimental research indicates that during this stage, the strong mossy-fiber inputs from the DG to CA3 dominate over the local recurrent-collateral inputs within CA3 [42, 59, 60]. This motivates the bipartite architecture of our model, where the internal CA3 connectivity is omitted.

According to the hippocampal index theory (HIT), neural units in the memory layer function as “indices” during early encoding to stabilize the memory patterns. HIT posits that each index binds all the feature neurons active in a memory so that, starting from partial cues (an incomplete set of active features), a complete memory can be retrieved by re-activating its index, a property known as pattern completion [39, 41, 53]. HIT also posits that each index selectively binds the features of only one memory, allowing pattern separation between different memories [38, 40–42, 54, 55]. However, it remains unproven whether the formation of neural indices with these properties is necessary to maximize the number of memories that can be stored in the network and the amount of information that can be encoded in the feature layer.

Here, we address this question by computing the memory and information capacity of the network with structural constraints defined in Fig. 4A. We set the number of units in the feature and memory layers to a 1/3 ratio, based on biological estimates of cell counts in CA3 and DG in rodents [61, 62]. The neuron coding levels are varied independently in the two layers. As a function of these coding levels, we compute the memory capacity, using Eqs. (7) and (3), and the information capacity, which is defined as the normalized entropy of possible states of feature neurons in the stored patterns (Methods). Convergent results from analytical and numerical estimates indicate that (a) memory capacity is maximized when the coding level is low or high in both layers (Fig. 4B, top); (b) information capacity is maximized when the coding level in the memory layer is low or high and the coding level in the feature layer is intermediate (Fig. 4B, middle); (c) memory and information capacity are more sensitive to the coding level in the memory layer compared to the feature layer; (d) as a consequence, to jointly maximize memory and information capacity (i.e., their product), the coding level must be low or high in the memory layer, whereas it is relatively unconstrained in the feature layer (Fig. 4B, bottom). Mathematically, the two optimal solutions are symmetric; however, biologically they are not: maximizing memory capacity through low or high coding levels corresponds to encoding information using the +1 (active) or -1 (silent) states, respectively. Thus, only the first solution (low coding levels) is biologically meaningful, since only active neurons can transmit information in the brain.

We then examine how the network configuration (set of *J*_*ij*_) that stores memory patterns at maximal capacity changes with varying coding levels in the memory and feature layers. Given a network configuration, we define an “Index Score” for each active memory unit in each stored pattern. This score quantifies how well the memory unit binds the feature units active in the same pattern while not binding the other units - in both directions, from the feature to the memory layer and vice versa (details in Methods). This is consistent with experimental evidence of bidirectional information flow between CA3 and DG [60, 63]. Fig. 4C shows two examples of memory units with high and low Index Scores, respectively. A high score means that the unit is activated by an incomplete set of features (a corrupted memory pattern) and, in turn, activates the full set of features of that memory (pattern completion) while inhibiting the features of other memories (pattern separation). A low score indicates that the network configuration doesn’t support this indexing function (i.e., conjunct pattern completion and separation) for that unit. We also define the Network Index Score as the mean index score of all active memory units across the stored patterns.

We find that the Network Index Score is maximized when the coding level in the memory layer is low, coinciding with the biologically meaningful condition that jointly optimizes memory and information capacity (Fig. 4D). In this condition, for a wide range of coding levels in the feature layer, the distributions of Index Scores for the memory units are entirely above zero, indicating a role in indexing the features of the respective memory patterns (Fig. 4DE, blue and red dots and distributions). Conversely, for other coding levels (for example, intermediate values in both layers), the distribution of Index Scores for the memory units is centered at zero, meaning that those units don’t support any indexing function (Fig. 4DE, yellow dot and distribution). Enforcing memory robustness (*κ >* 0) does not change these results (Supplemental Fig. 5).

In conclusion, maximizing memory and information capacity in two-layer networks, such as the one shown in Fig. 4A, requires coding levels and synaptic weight configurations that support an indexing function for the memory layer, as posited by the HIT. An important factor in the model architecture of Fig. 4A is that the memory layer has more neurons than the feature layer. If this is reversed, the memory capacity becomes more sensitive to the coding level in the feature layer, and the conditions that jointly maximize memory and information capacity no longer overlap with those of maximum Network Index Score (Supplemental Fig. 6). This result supports the hypothesis that memory indices are stored in brain areas that have more neurons than the regions storing the memory features. Thus, our result argues against an indexing scheme where the hippocampus directly indexes entire cortical (feature) modules. Instead, it supports a hierarchical scheme, where the DG would store high-order indices (i.e., indices of indices) and CA3 would store the features of those indices, which in turn may be part of a network that forms lower-order indices of features stored in cortical modules [38–40, 54, 56, 64].

## III. DISCUSSION

The first contribution of this research is the calculation of perceptron capacity for a broad class of binary classification problems, where the input patterns and output labels can have arbitrary statistics, provided that the input states have finite means and variances, and are either independent or linearly correlated. We found that the perceptron capacity depends only on the mean value of the labels (*m*^*out*^). Several previous studies used similar statistical mechanics techniques - pioneered by Gardner [16] - to estimate the perceptron capacity for different classification scenarios or under different constraints [26, 27, 30, 31, 34, 65]. For example, in [26], the capacity of perceptrons was obtained for higher-order classification problems, where a mapping is learned between groups of inputs and groups of labels. In [30, 31], the capacity was calculated for classifying manifold objects. In [27, 34, 65] the authors addressed classification problems with correlated input/input or input/output states, and unbiased outputs (*m*^*out*^ = 0). A consistent result for the upper bound, *α*_*p*_ = 2, was also obtained using a different geometrical approach, based on the assumption that the patterns are in a general position, i.e., no subset of *N* or fewer patterns are linearly dependent [25, 66]. However, this approach does not apply to cases with biased outputs (*m*^*out*^ ≠ 0) and finite classification stability (*κ* ≠ 0). In summary, to the best of our knowledge, this is the first study to provide an explicit formula for the perceptron capacity in the case of arbitrary distributions of input states and both biased and unbiased outputs (any *m*^*out*^). This result served as the foundation for deriving a general analytical formula for the memory capacity of networks with varying coding levels.

We thus computed the capacity of Hopfield models with heterogeneous coding levels and arbitrary architectures. This result extends previous literature, which addressed the case of homogeneously connected (dense or diluted) Hopfield networks, with continuous or discrete synapses, and binary or threshold-linear units [6, 16–22, 67]. The case of heterogeneous networks was previously addressed in [6, 23, 24], but only for graphs with symmetric weights (learned by symmetric Hebbian learning) and a constant coding level across all neurons. Our mathematical formula (7) shows that the network capacity is a function of both the coding level and the number of inward (but not outward) connections of each cell.

While we considered networks with (−1, 1) units, our results generalize to (0, 1) units. Indeed, for a perceptron, replacing the −1 labels with 0 does not affect linear separability of the patterns, and changing the input states in a similar way only alters the input-state distributions, which do not affect the capacity. The capacity of a Hopfield model is defined as the minimum capacity of the perceptrons in the network, and is therefore also invariant to this transformation. The formula (7) also implies that in two-layer networks with only inter-layer connectivity the capacity must be linear in the number of units of one of the layers (specifically, the input layer of the perceptron with minimum capacity). This is consistent with the scaling of memory capacity found for Modern Hopfield networks [8, 9]. Altogether, our analytical results enable the calculation of memory capacity for a broader class of networks than previously possible. In this study, we examined the capacity of two biologically inspired types of networks: a general class of heterogeneous networks and a class of two-layer memory networks.

In the first case, we assessed how the amount of heterogeneity in the networks - specifically, the variability of coding levels and number of inward connections (in-degrees) across neurons - affects memory capacity. We found that, in general, memory capacity decreases with increasing heterogeneity in either parameter. This result is consistent with previous findings that maximizing memory capacity in networks with positive weights and homogeneous coding levels leads to a smaller variance in in-degrees compared to out-degrees [7]. It also aligns with the known decrease in memory capacity as correlations among the patterns increase [32], since coding levels different from *p*_*i*_ = 0.5 lead to correlated patterns. We also found that the reduction in memory capacity caused by heterogeneity is mitigated under specific conditions defined by the joint statistics of neuron coding levels and in-degrees. In particular, in a regime of low-coding levels (*p*_*i*_ *<* 0.5), which is more energy efficient and biologically plausible [45–50], coding levels and in-degrees should be positively correlated (i.e., neurons with fewer inward connections should have lower coding levels) to maximize the network memory capacity. This result represents a normative prediction for real brain networks, which could be tested experimentally. While direct measurements of synaptic connectivity are generally unfeasible *in vivo*, several methods exist to infer functional or statistical interactions between cells from spike data. Some of these methods [68–70] yield sparse interaction graphs by carefully avoiding overfitting and could thus be used to estimate neuron in-degrees.

We then considered networks composed of two reciprocally (but not internally) connected layers, similar to the BAM model but with potentially asymmetric synaptic weights [35]. The two layers differ in the number of neurons and coding levels. We found that the memory capacity of this class of networks is maximized when the layer with more neurons (which we call the memory layer) has a low coding level, i.e., sparse neuronal activation. Under this condition, neurons active in the same layer have an indexing function, binding the memory features active in the other layer, as hypothesized by HIT and in agreement with most recent “cognitive map” theories [71–74]. Meanwhile, the information capacity of the feature layer is also maximized for this network configuration. Interestingly, the DG and CA3 regions of the hippocampus seem to have these properties: the DG has 3 to 5 times more neurons than CA3, based on biological estimates in rodents [61, 62], and very sparse coding levels [75]. This correspondence suggests that the DG may store indices of memories initially encoded in CA3, supporting both pattern completion and pattern separation in the latter [39, 41, 53, 76, 77]. Our results argue against an indexing scheme where neurons in the hippocampus directly bind features in the cortex. In networks where the feature layer has more neurons than the memory layer (as suggested by a cortical-hippocampal model), we found that maximizing the indexing function of the memory layer does not maximize memory capacity. Instead, our results support a hierarchical indexing scheme, where the DG binds features in CA3, which may be part of a broader network that binds features in the cortex [38]. Additionally, the indexing function of the DG may be supported by adult neurogenesis of granule cells, which provides a mechanism for updating indices as new memories are continually formed [76–80].

## MATERIALS AND METHODS

### A. Numerical estimation of the perceptron capacity

We generate *P* random pattern-label pairs and train the perceptron to store the pairs using the “sklearn.svm.LinearSVC” function in the scikit-learn package [81], which implements the support vector machine (SVM) algorithm for pattern classification. We use the binary search algorithm to find the maximum number of pattern-label pairs *P* that the perceptron can store with an error rate of 0.01 or less. We then compute the mean and standard deviation of the maximum number of patterns over 10 iterations of this process. The number of perceptron input units used in the simulations is 1000.

The analytical capacity is given by solving Eqs. (3) with numerical methods provided by the “scipy.optimize.root” function [82].

### B. Gaussian Markov sampling

For the perceptron with correlated inputs (Fig. 1GH), we sample the input states from a Gaussian Markov chain according to the following formula:

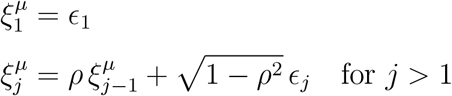

where each *ϵ*_*j*_ is drawn from a standard normal distribution.

### C. Numerical estimation of the capacity of the Hopfield model

The Hopfield model is formally equivalent to *N* independent perceptrons. Each perceptron *i* is trained to associate label 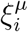 to pattern 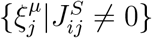 using the “sklearn.svm.LinearSVC” function, where 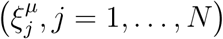 is a memory pattern. We use the binary search algorithm to find the maximum number of memory patterns that the network can store with an error rate of 0.01 or less. The error rate is defined as the mean fraction of perceptrons per pattern that don’t satisfy all the classification constraints. We then compute the mean and standard deviation of the maximum number of patterns over 10 iterations of this process.

The numbers of units used in simulations of the two-layer network of Fig. 4 are *N*_1_ = 150 for the feature layer and *N*_2_ = 450 for the memory layer. The number of units used in simulations of the randomly connected heterogeneous networks of Fig. 3 is *N* = 500.

### D. Index Score

For the two-layer network of Fig. 4, we denote the portions of each pattern stored in the feature and in the memory layer as ***ξ***^**1**,***µ***^ and ***ξ***^**2**,***µ***^, respectively. The numbers of units in the feature and in the memory layer are denoted as *N*_1_ and *N*_2_, respectively.

We binarize the inward and outward weights of unit *i* in the memory layer using the formulas 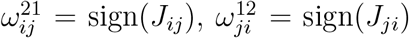, so that 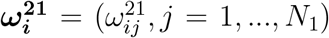 and 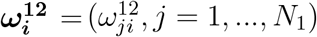 are the binarized weight vectors from the feature layer to memory unit *i* and from memory unit *i* to the feature layer, respectively.

For each stored pattern (***ξ***^**1**,***µ***^, ***ξ***^**2**,***µ***^) and each unit in the memory layer that is active in that pattern, we compute the average cosine similarity between the binarized weight vectors and the portion of the pattern in the feature layer:

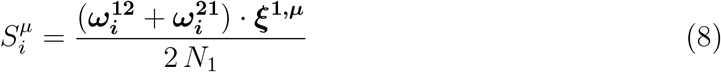

We also calculate the baseline (control) cosine similarity as the expected value of 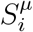 random patterns ***ξ***^**1**,***µ***^ (with the same neuron coding level *p*_1_):

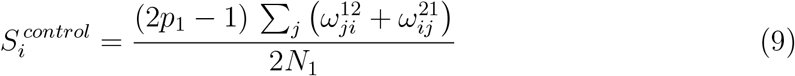

We define the Index Score of each neuron active in the memory layer in each stored pattern as

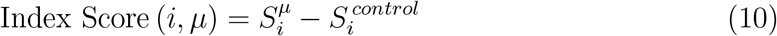

We define the Network Index Score as

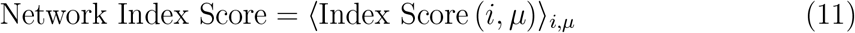

### E. Information capacity in the feature layer

We denote the coding level in the feature layer by *p*_1_ and the neuron numbers in the feature and the memory layer by *N*_1_, *N*_2_, respectively.

The information capacity (per synapse) is calculated as the entropy of the states of feature neurons across all memory patterns, normalized by the total number of possible synapses in a network of size *N*_1_ + *N*_2_:

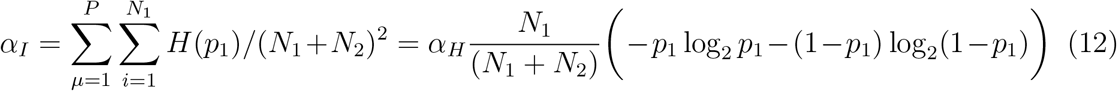

### F. Generation of heterogeneous networks

To generate the structure *J*^*S*^ of the randomly connected heterogeneous networks shown in Fig. 3, we use the following procedure: we assign an inward-connection probability 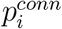 to each neuron *i*; 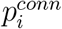 is sampled from a uniform distribution with mean 0.5; we then set 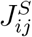 to 1 with probability 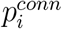 and to 0 with probability 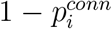, for every *j*. We also assign a coding level *p*_*i*_ to each neuron *i*; *p*_*i*_ is sampled from another uniform distribution with mean 0.25. To generate networks with different levels of heterogeneity, we vary the standard deviations of the distributions of 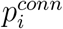 and *p*_*i*_, while keeping their means constant.

To enforce a positive correlation between coding levels and in-degrees, we permute the coding levels *p*_*i*_ so that the neuron with the lowest *p*_*i*_ is associated with the lowest 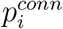, the neuron with the second lowest *p*_*i*_ is associated with the second lowest 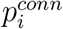, and so on. We use the inverse procedure (associating low *p*_*i*_ with high 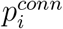) to enforce a negative correlation between coding levels and in-degrees.

## ACKNOWLEDGMENTS

The authors thank Edward Han for fruitful discussions on the neural substrates of the hippocampal index theory, and Rémi Monasson, L-ukasz Kuśmierz, Naoki Hiratani, and Ilya Monosov for useful comments on the manuscript. G.T. acknowledges support from the Sloan Foundation under Grant No. FG-2024-22163.

## APPENDIX A: ANALYTICAL CALCULATION OF THE PERCEPTRON CAPACITY

We present here the complete calculation of the maximal capacity of a perceptron with arbitrary (independent) distributions of input states and output labels.

### Definition of the problem

We define *P* input patterns 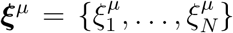, where *µ* = 1, …, *P* indexes the pattern and each input state 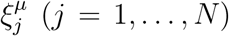 is sampled from an arbitrary distribution with finite mean *m*_*j*_ and variance 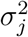. We do not impose any constraint on these distributions insofar as the states of different neurons are statistically independent. Each pattern is associated with a label *χ*^*µ*^. The labels are drawn from a Bernoulli distribution with mean *m*^*out*^: *p*(*χ*^*µ*^ = 1) = (1 + *m*^*out*^)*/*2 and *p*(*χ*^*µ*^ = −1) = (1 − *m*^*out*^)*/*2.

We denote the normalized capacity *α*_*p*_ = *C*_*p*_*/N*, where *C*_*p*_ is the maximal number of pattern-label pairs that the perceptron can learn to classify, according to the classification constraints:

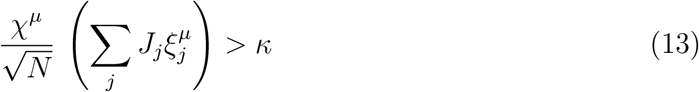

for every input pattern indexed by *µ*; *κ* ≥ 0 is the stability parameter (Fig. 1B).

To calculate *α*_*p*_, we define the following elliptic regularization on the perceptron synaptic weights {*J*_*j*_}:

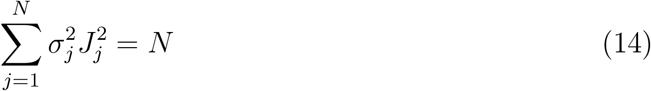

This constraint can be viewed as a normalization of the postsynaptic input ranges. When all inputs have the same variance, Eq. (14) reduces to the standard spherical regularization used by Gardner [16]. As discussed in Appendices B and C, the difference between our elliptic and Gardner’s spherical constraint is relevant only when the stability parameter *κ* in Eq. (13) is strictly positive. When *κ* = 0, any solution that satisfies (14) can be mapped to exactly one solution that satisfies a spherical constraint, or any other elliptic constraint with coefficients different from 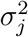. Thus, when *κ* = 0, the maximal capacity is invariant to changes in the coefficients of constraint (14).

Following the method introduced by Gardner [16], we define the fractional volume of the weights that satisfy both conditions (13) and (14) in the space of weights constrained by (14):

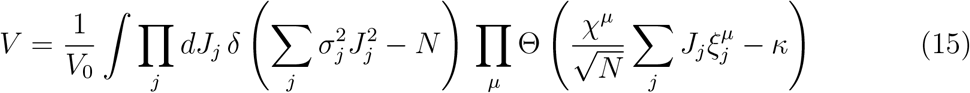

where

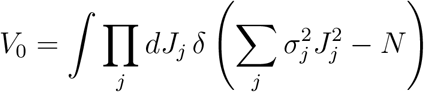

*V* represents the volume of the solutions for a given realization of the pattern-label pairs. As the number of patterns *P* increases, *V* diminishes. The capacity *C*_*p*_ = *α*_*p*_ *N* is defined as the number of patterns at which ⟨log *V* ⟩_*ξ,χ*_ → −∞, where ⟨·⟩_*ξ,χ*_ denotes the average over all possible realizations of input states 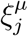 and output labels *χ*^*µ*^. Above this limit, for a set of realizations with non-zero measure, no {*J*_*j*_} configuration can satisfy all the constraints (13) (and vice versa, below this limit, the perceptron can store all the patterns with probability 1).

In practice, the calculation of ⟨log *V* ⟩_*ξ,χ*_ is done using the replica method: *n* replicas of the perceptron are introduced and the typical volume of the solutions for the *n* replicas ⟨*V* ^*n*^⟩_*ξ,χ*_ is obtained using the saddle-point method for *N* → ∞, and in the limit *n* → 0 (assuming analytic continuation of the volume from positive integers). Finally, the capacity *α*_*p*_ is determined by solving for ⟨*V* ^*n*→0^⟩_*ξ,χ*_ → 0, so that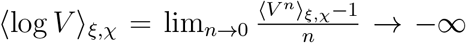. All the steps are recapitulated in detail below.

#### Calculation of the volume of the solutions for *n* replicas

The volume of the solutions for the *n* replicas (indexed by the superscript *α*) is defined as:

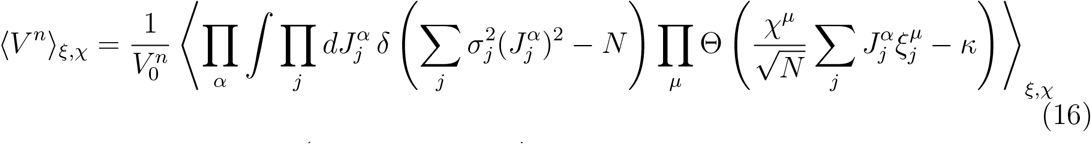

with 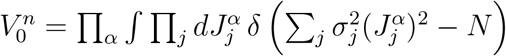.

We substitute the Heaviside step function Θ with its integral representation:

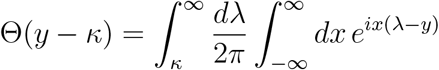

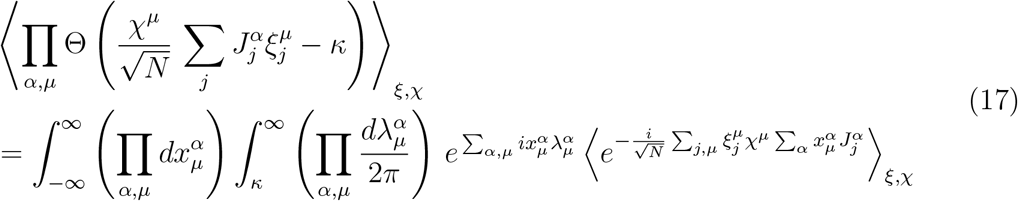

By taking the average over the distributions of 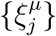 we obtain:

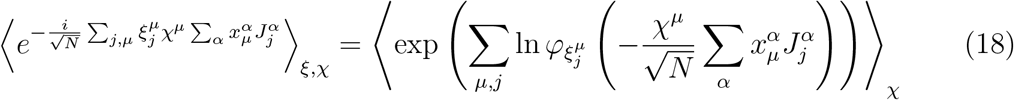

where 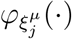 is the characteristic function of 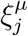.

Expanding 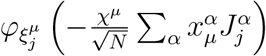 and its logarithm to second order around zero and using the constraint (14), we obtain:

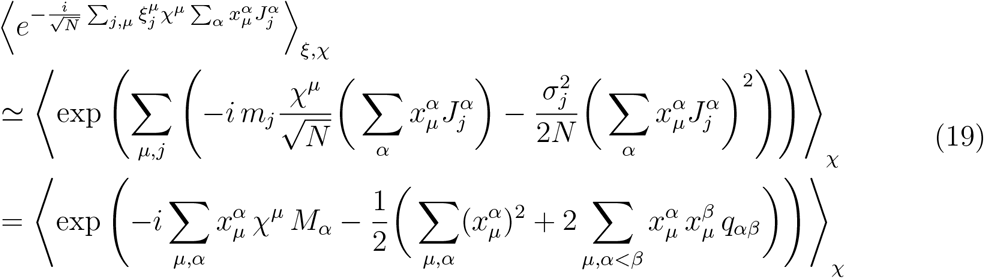

where we have introduced the variables

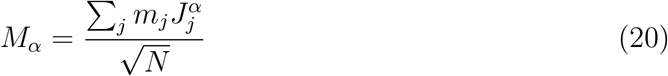

and

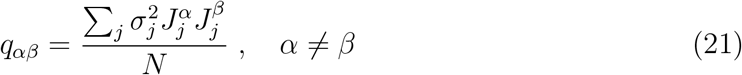

Substituting (19) back into (17):

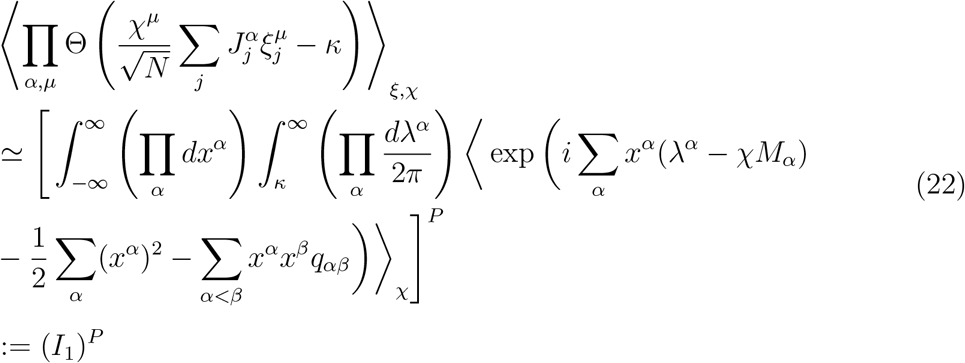

For convenience, we have introduced the symbol *I*_1_ to denote the integral. Using (20) and (21), we can rewrite ⟨*V* ^*n*^⟩_*ξ,χ*_ with *M*_*α*_ and *q*_*αβ*_ as integral variables:

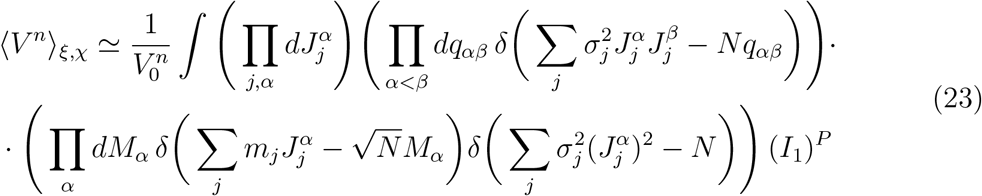

We now use the integral representation of the *δ* function

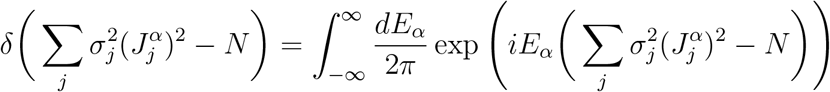

We do the same for the other *δ* functions in Eq. (23), introducing two additional variables *F*_*αβ*_ and *K*_*α*_:

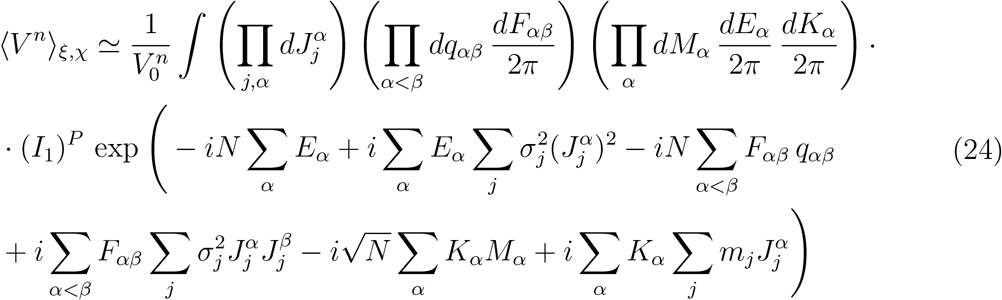

Eq. (24) can be expressed as:

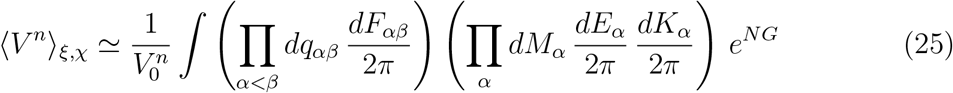

where

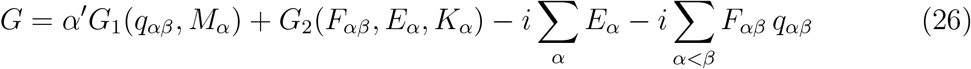

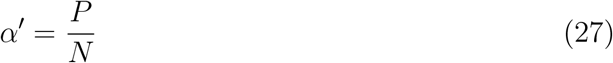

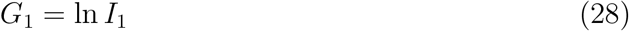

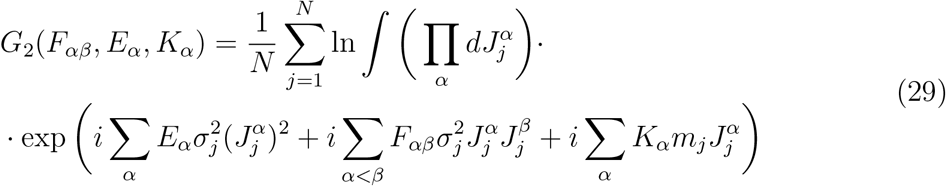

As *N* → ∞ and *P* → ∞ with *α*^′^ = *P/N* finite, 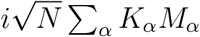 in Eq. (24) is small compared to the other terms and is discarded.

Below we estimate *G*_1_, *G*_2_, and the volume, Eq. (25), using the saddle-point method as *N* → ∞ (*α*^′^ finite). To obtain the saddle-point equations, we take the limit *n* → 0 (assuming analytic continuation of ⟨*V* ^*n*^⟩_*ξ,χ*_ from positive integers) and we use the replica-symmetric ansatz [16]:

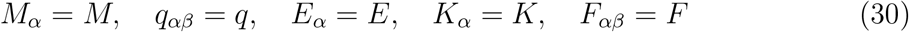

#### Calculation of *G*_1_

Using the replica symmetric ansatz, *I*_1_ simplifies to:

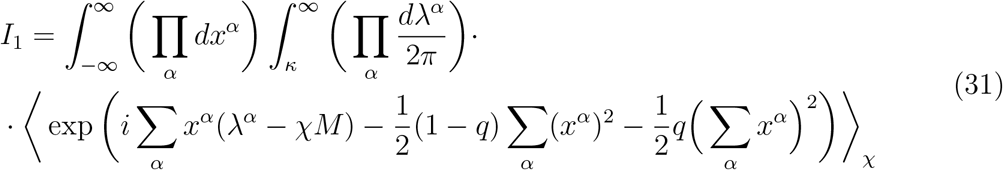

The term containing 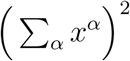 is replaced by a Gaussian integral, so that *I*_1_ factorizes over the replicas *α*. Defining 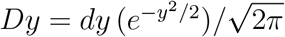, we have:

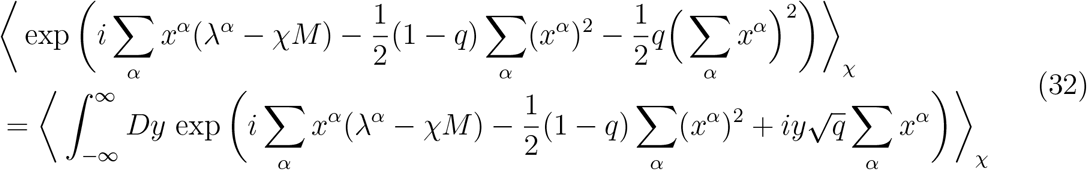

Thus:

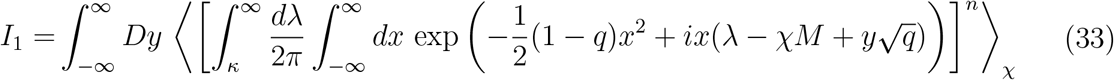

We introduce the following notation for convenience

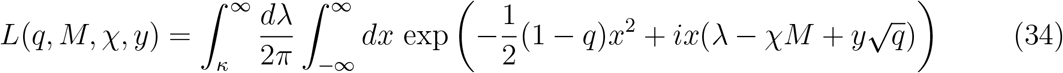

so that

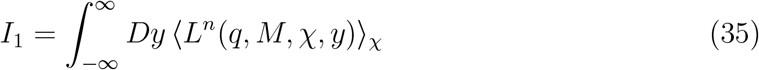

Solving the Gaussian integral in *x* in Eq. (34), we obtain:

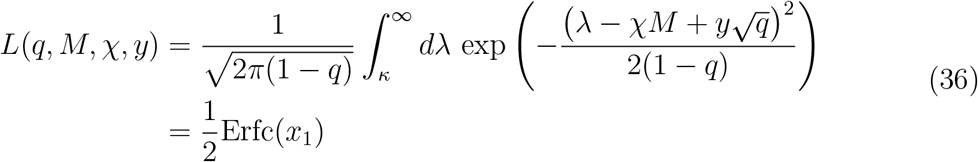

with Erfc 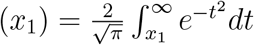 and 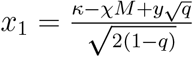.

We now take the limit of *G*_1_ as *n* → 0:

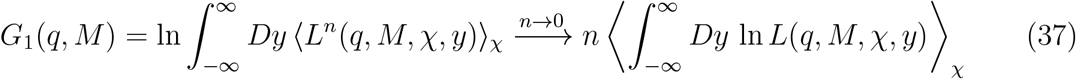

where we have used 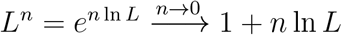.

Finally we write the average over the *χ* values explicitly:

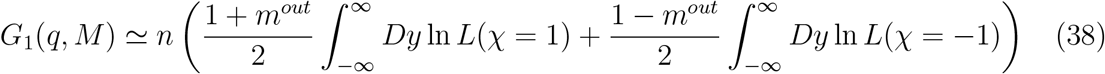

where *L*(*χ* = ±1) is the simplified notation for *L*(*q, M, χ* = ±1, *y*).

#### Calculation of *G*_2_

Using the replica symmetric ansatz, Eqs. (30), *G*_2_ simplifies to:

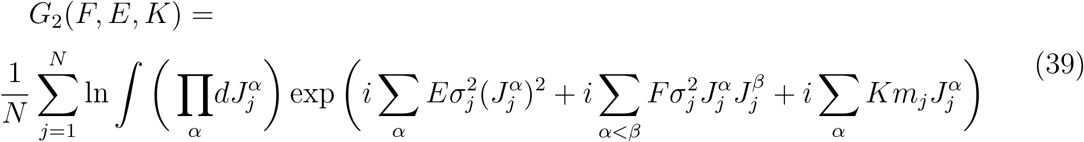

We define two statistics 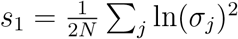 and 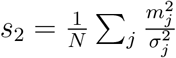 related to the distributions of input states, and solve the multidimensional Gaussian integral in Eq. (39):

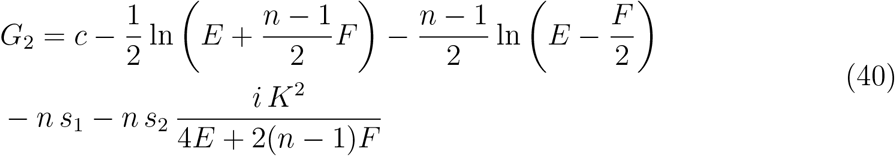

where *c* is a constant, independent of the relevant variables.

We take the limit *n* → 0:

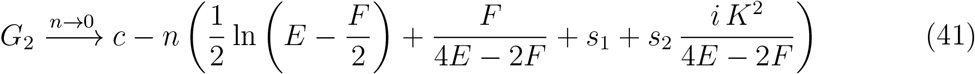

### Derivation of the saddle-point equations

We substitute the results obtained in the limits *N* → ∞, *n* → 0 into Eq. (26):

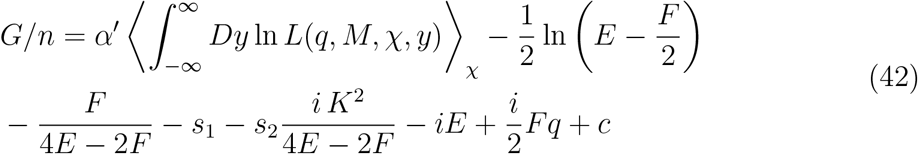

By the method of steepest descent, the volume is estimated at the saddle-point:

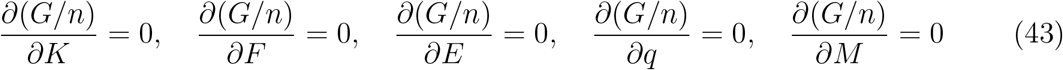

The first three equations in (43) give *K, F, E* as functions of *q*:

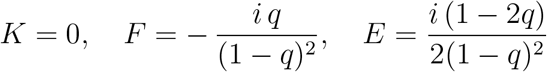

Substituting them back into (42) we have:

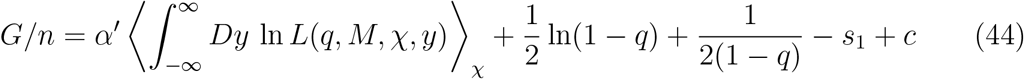

As *α*^′^ increases, the typical volume ⟨*V* ^*n*→0^⟩_*ξ,χ*_ decreases and the solutions 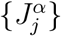 become more correlated. Approaching the maximal capacity (*α*^′^ → *α*_*p*_), the typical volume shrinks to 0 and the solutions 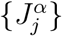 for all the replicas converge to the same configuration. It follows that 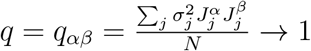 for all *α, β*, by constraint (14).

Thus, to estimate the maximal capacity *α*_*p*_, it is sufficient to solve the two remaining saddle-point equations in (43) for *q* → 1:

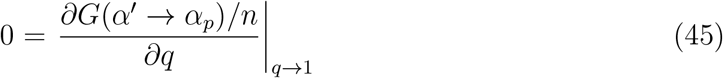

and

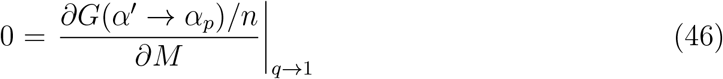

### Solving Eqs. (45) and (46)

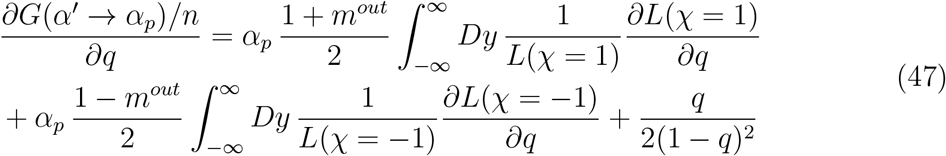

For the first term on the right-hand side of Eq. (47), substituting the expression for *L*(*χ* = 1) from (36), we obtain:

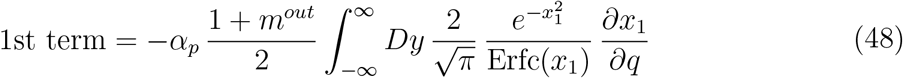

with 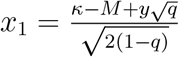.

Approximating the complementary error function 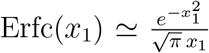 for *x*_1_ ≫ 0 [83], and taking the limit *q* → 1, the integration (48) simplifies to:

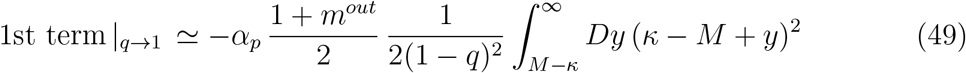

For the second term on the right-hand side of Eq. (47), a similar approach yields:

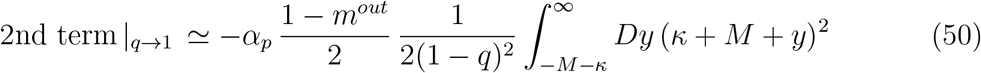

Substituting into Eq. (45), we obtain:

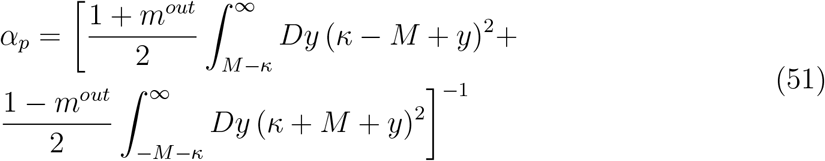

Following the same procedure for Eq. (46), we obtain the equation that gives the value of *M* :

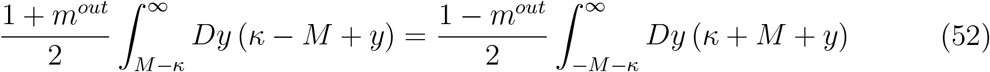

Eqs. (51) and (52) together provide the analytical solution for the maximal capacity of the perceptron when *m*_*j*_ ?= 0 for at least one input unit. When *m*_*j*_ = 0 for all input units, *M* = 0 by definition. Thus, Eq. (46) is not defined and the maximal capacity is given by Eq. (51) with *M* = 0:

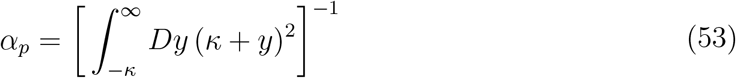

*Special case: κ* = 0

When the stability parameter *κ* = 0 (such that condition (13) is sufficient for the input patterns to be stored as fixed points in the network but not sufficient for the basins of attraction to be finite), the formula for the maximal capacity further simplifies. Integrating by parts in Eq. (51), solving the integrals in Eq. (52), and substituting the resulting equality into Eq. (51), we obtain:

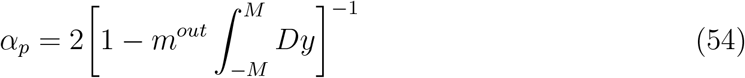

If *m*_*j*_ = 0 for all input units, then *M* = 0 and *α*_*p*_ = 2; otherwise, *M* is determined by:

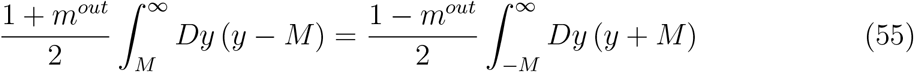

We prove below that when *κ* = 0 the maximal capacity is independent of the elliptic regularization (14).

## APPENDIX B: INDEPENDENCE OF THE CAPACITY FROM THE ELLIPTIC REGULARIZATION WHEN *κ* = 0

We demonstrate that there exists a bijective relation between the sets of solutions {*J*_*ij*_} that satisfy (13) with *κ* = 0 under two arbitrary (different) elliptic constraints: constraint A, Σ_*j*_ *a*_*j*_*J*_*j*_^2^ = *N*, and constraint B, Σ_*j*_ *b*_*j*_*J*_*j*_^2^ = *N*.

Given input patterns 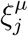 and their corresponding labels *χ*^*µ*^, for every set 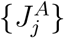 that satisfies constraint A and the classification conditions

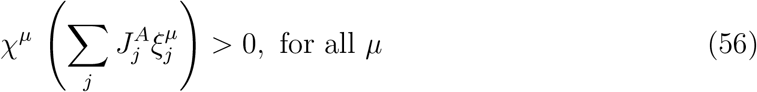

there exists a set 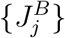 defined as

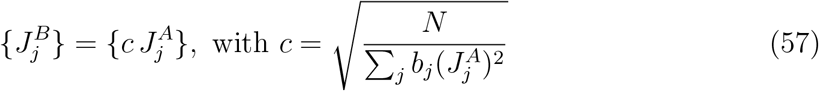

that satisfies constraint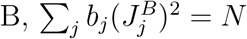, and the classification conditions:

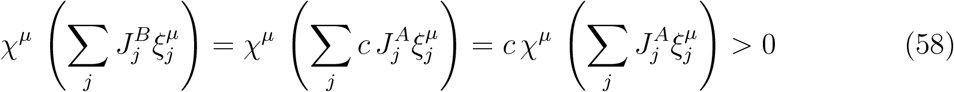

Similarly, for every set 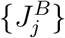that is a solution under constraint B, there exists a set 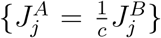 that is a solution under constraint A. In conclusion, there exists a one-to-one mapping between the solution spaces under constraints A and B. Since A and B can be chosen arbitrarily, this bijection implies that the maximal capacity, for *κ* = 0, is not affected by the specific elliptic constraint used in the calculation.

## APPENDIX C: COMPARISON WITH GARDNER’S SOLUTION

In Gardner’s calculation [16, 17], all units have binary, independent, identically distributed states, with *m*^*out*^ = *m*_*j*_ = *m* and 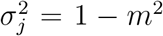. This condition corresponds to a special case within our framework.

To address the case of arbitrary {*m*_*j*_} and 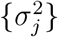, we implemented some modifications in our calculation. These modifications make use of an elliptic constraint on the weights, which is effectively equivalent to an input-range normalization:

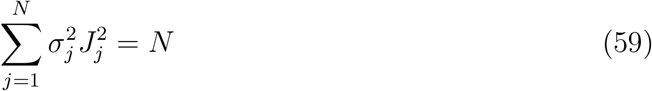

This is in general different from Gardner’s spherical constraint 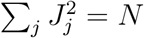.

In the special case of binary, independent, identically distributed states with mean *m* and variance 1−*m*^2^, our solution for the maximal capacity reduces to Gardner’s solution upon a rescaling of the stability parameter *κ*. In particular, the elliptic constraint simplifies to a spherical form, 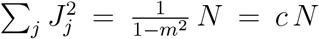, which differs from Gardner’s constraint 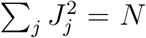 by the coefficient *c*. As previously shown in [27], the effect of the coefficient *c* = 1*/*(1 − *m*^2^) is equivalent to a reduction in the stability *κ* of the patterns by a factor 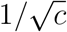. Thus, expressing the capacity *α*_*p*_ under the spherical constraint 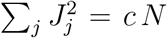 as *α*_*p*_(*κ, m, c*), the following equivalence holds [27]:

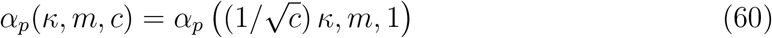

Indeed, substituting 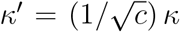 into Eqs. (51) and (52), and defining 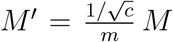, we retrieve Gardner’s solution for the maximal capacity:

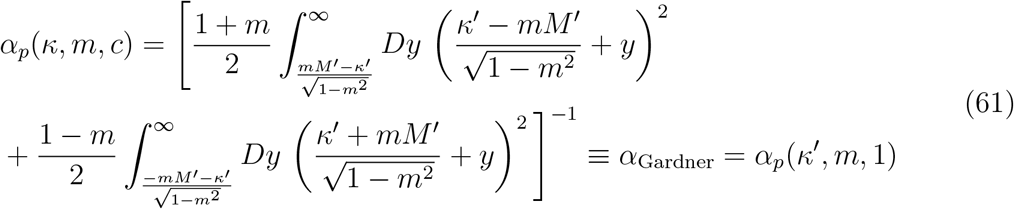

where *M* ^′^ is given by

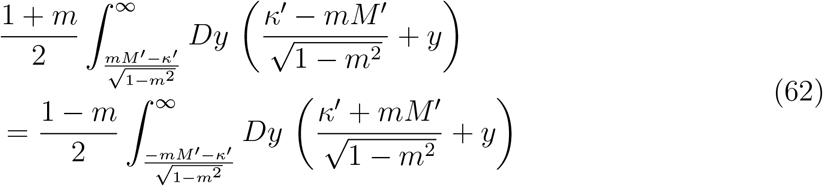

