## Supplemental Figures for "Maximizing memory capacity in heterogeneous networks"

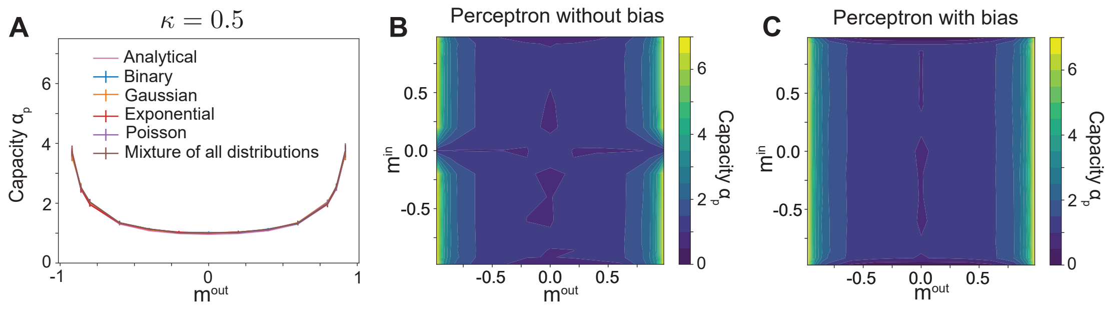


Figure 1: **The capacity of a perceptron with finite stability ($\kappa > 0$)**. Same analyses as in Fig. 1CDE after replacing $\kappa = 0$ with $\kappa = 0.5$. **A**. Maximal capacity of a perceptron as a function of the output mean $m^{out}$; analytical solution, Eqs. (51-52) in Appendix A, and mean ± std over 10 numerical estimates (N=1000, Methods) for perceptrons with bias and different input distributions (legend). The capacity is lower for $\kappa > 0$ than for $\kappa = 0$ but follows the same trend (compare with Fig. 1C). **B**. Numerical estimates of the maximal capacity of a perceptron without bias. Input (output) states are independently sampled from binary distributions with mean $m^{in}$ ($m^{out}$). The capacity varies as a function of $m^{out}$ but not $m^{in}$, provided $m^{in} \neq 0$ (with slight modulations due to finite-size effects), as in Fig. 1D. When $m^{in} = 0$, the analytical solution for the capacity is given by Eq. (53), Appendix A. **C**. Same as (B) for a perceptron with bias. The capacity depends solely on $m^{out}$, regardless of whether $m^{in} = 0$ or $m^{in} \neq 0$, as in Fig. 1E.

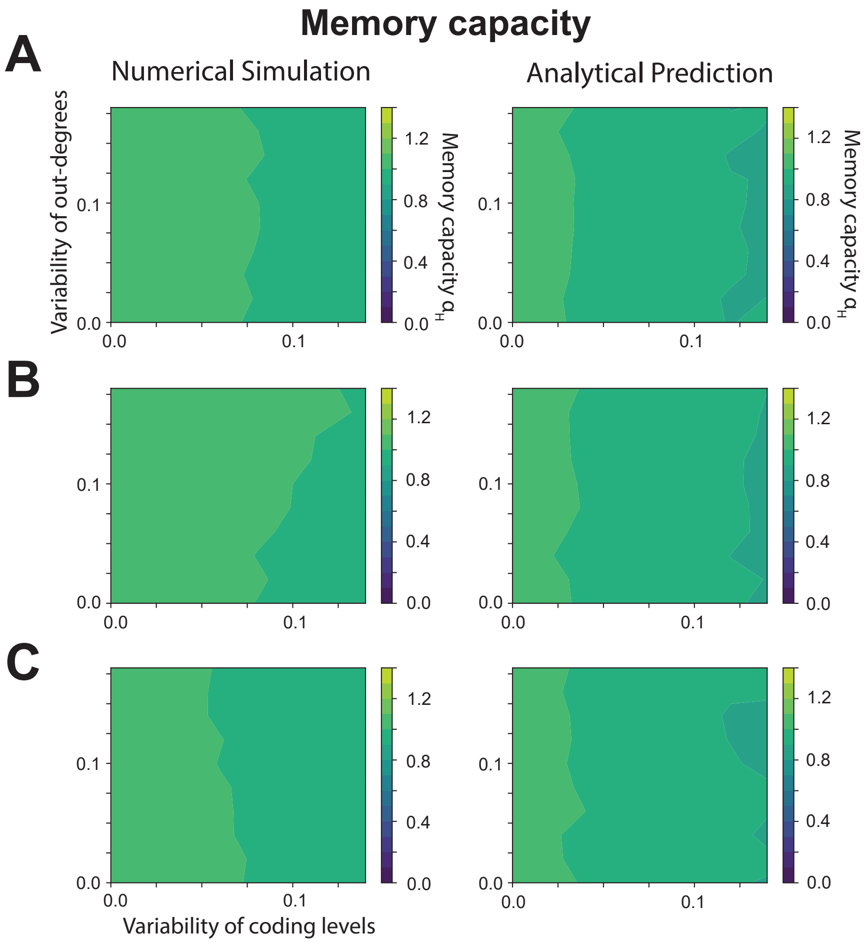


Figure 2: **Memory capacity is unaffected by the out-degrees of neurons in the network**. Numerical (left) and analytical (right) estimates of memory capacity are shown (on the same scales as in Fig. 3EFG for comparison). Here, the in-degrees are independent of the coding levels, with the inward-connection probability set to 0.5 for every cell. The outward-connection probability for each cell is sampled from a uniform distribution with a mean of 0.5 and a variable standard deviation ($y$-axis). The coding levels are sampled from a uniform distribution with a mean of 0.25 and a variable standard deviation ($x$-axis). The capacity slightly decreases with increasing variability of coding levels, as expected from Fig. 3E, but it is not affected by the variability of out-degrees, regardless of whether out-degrees and coding levels are (**A**) independent, (**B**) positively correlated, or (**C**) negatively correlated. Minimal variations along the $y$-dimension are due to numerical noise.

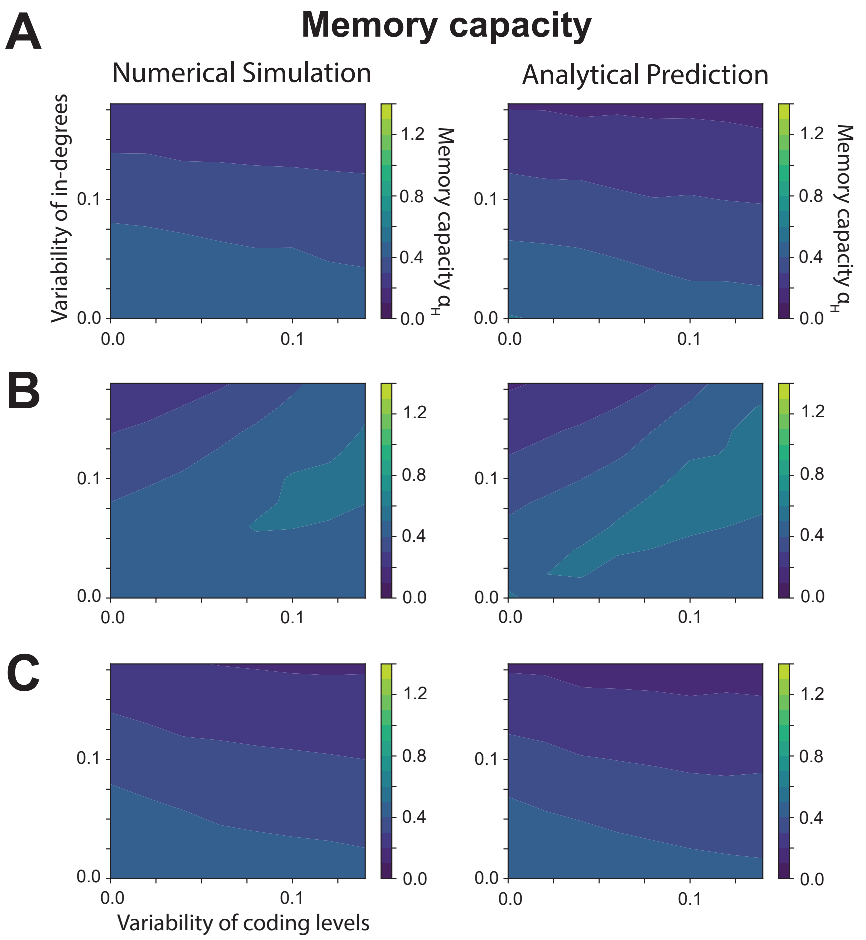


Figure 3: **Memory capacity of heterogeneous random networks with finite stability ($\kappa > 0$)**. Numerical (left) and analytical (right) estimates of memory capacity are shown for $\kappa = 0.5$ (on the same scales as in Fig. 3EFG for comparison). Coding-level and in-degree distributions are defined as in Fig. 3 and Methods. **A**. When in-degrees and coding levels are independent, the capacity decreases as the variability of either parameter increases. **B**. When in-degrees and coding levels are positively correlated, the capacity of heterogeneous networks (with high variability in both parameters) is higher than when these parameters are independent and is similar to the capacity of homogeneous networks (with low variability). **C**. When in-degrees and coding levels are negatively correlated, the capacity decreases faster with increasing variability than when the two parameters are independent. While the capacity is consistently lower when $\kappa > 0$ than when $\kappa = 0$, the variability of in-degrees and coding levels, along with their joint statistic, affects memory capacity in similar ways (compare with Fig. 3EFG, respectively).

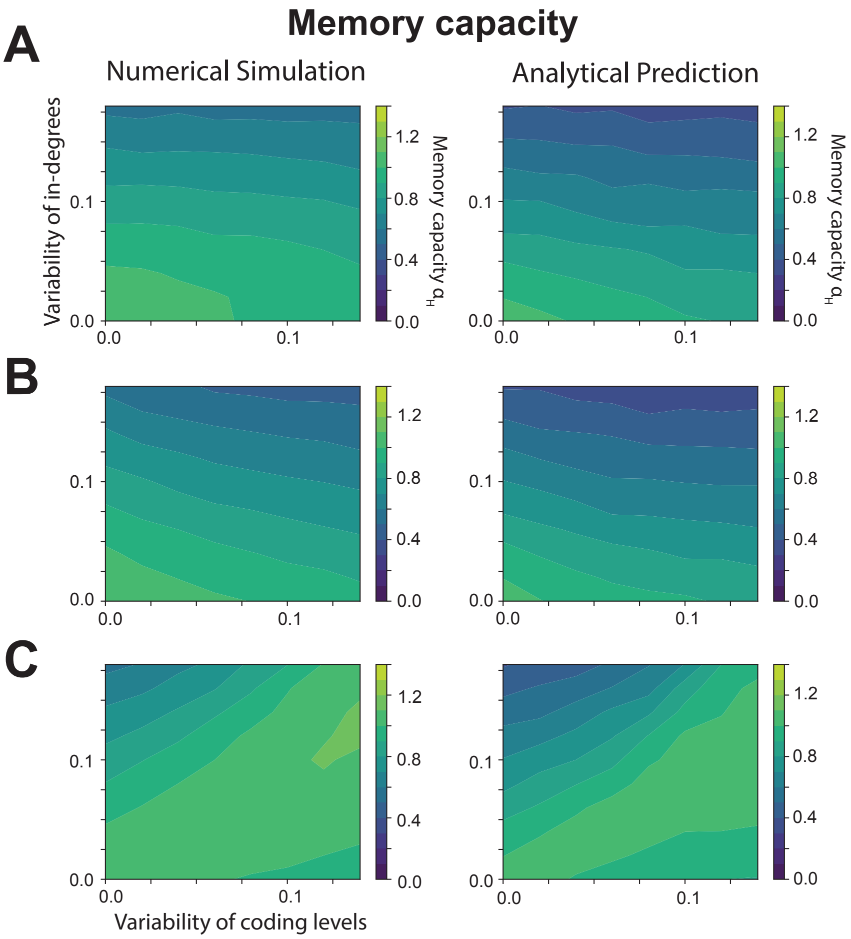


Figure 4: **Memory capacity of heterogeneous random networks in the regime of high coding levels ($\mathbf{p_i > 0.5}$ for all neurons)**. Coding levels are sampled from a uniform distribution with a mean of 0.75, while inward-connection probabilities are sampled from a uniform distribution with a mean of 0.5. In-degrees and coding levels are (**A**) independent, (**B**) positively correlated, or (**C**) negatively correlated. The results are reversed compared to the low coding-level regime (shown in Fig. 3EFG): the memory capacity of heterogeneous networks with high coding levels is maximized when in-degrees and coding levels are negatively correlated (C).

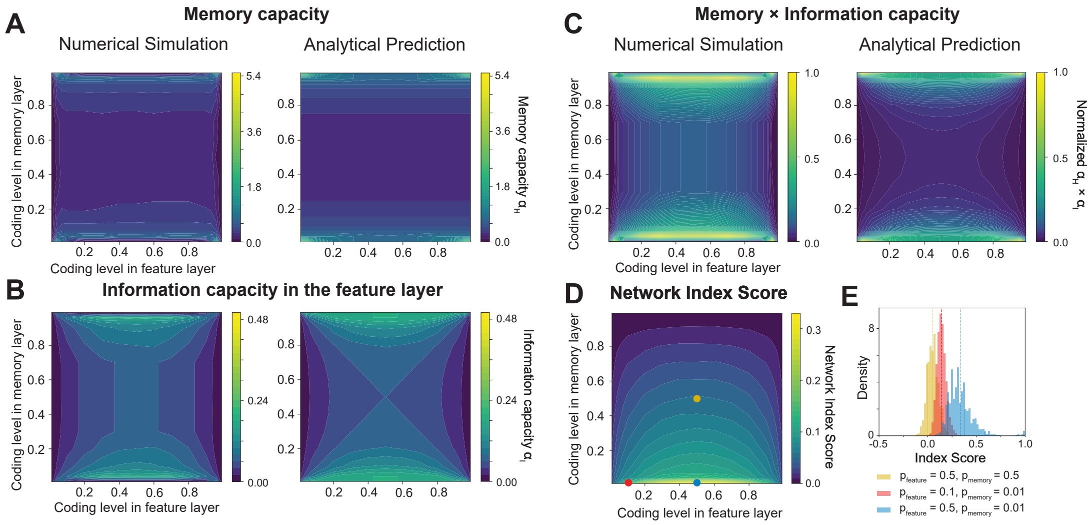


Figure 5: **Capacity of two-layer networks with finite stability ($\kappa > 0$) and alignment with hippocampal index theory (HIT).** Results are shown for the two-layer network depicted in Fig. 4A, with stability $\kappa = 0.5$. Memory (**A**), information (**B**), and memory $\times$ information (**C**) capacity (the latter normalized by the maximum value) as a function of the coding levels in the feature and memory layers: numerical (left) and analytical (right) estimates are shown on the same scales as in Fig. 4B for comparison. Both memory and information capacities are more sensitive to the coding level in the memory layer. When this is low or high, both capacities are maximized, as seen in the network with $\kappa = 0$ (compare with Fig. 4B). **D**. The Network Index Score varies with coding levels in the feature and memory layers. It is maximum when the coding level in the memory layer is low, which overlaps with one of the conditions of maximum memory and information capacity, as seen in the network with $\kappa = 0$ (compare with Fig. 4D). **E**. Distributions of Index Scores for the active memory units in the stored patterns are shown for different coding levels: both intermediate in the memory and feature layers (yellow), low and intermediate, respectively (blue), or both low (red), corresponding to the same-colored dots in (D). When the coding level in the memory layer is low, the distributions of Index Scores are entirely above zero, indicating an indexing function for the memory units, as posited by HIT. Altogether, enforcing memory robustness ($\kappa > 0$) does not qualitatively change the results of Fig. 4.

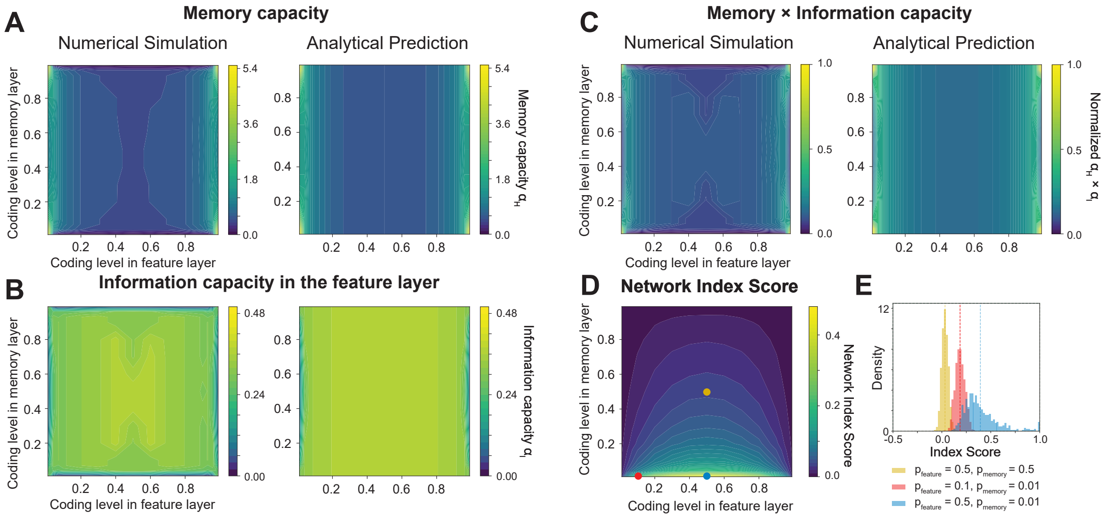


Figure 6: **Capacity and Index Scores of two-layer networks with more neurons in the feature layer than in the memory layer**. Results are shown for the two-layer network depicted in Fig. 4A, with an inverted ratio (3 instead of 1/3) between the number of neurons in the two layers. Memory (**A**), information (**B**), and memory $\times$ information (**C**) capacity (the latter normalized by the maximum value) as a function of the coding levels in the feature and memory layers: numerical (left) and analytical (right) estimates. Both memory and information capacities are more sensitive to the coding level in the feature layer, unlike in Fig. 4B. Note that information capacity is low for extreme coding levels in the feature layer (either very low or very high), whereas it is relatively high and uniform for intermediate coding levels (B). **D-E**. The Index Scores vary with coding levels in the feature and memory layers in a manner similar to the network analysed in Fig. 4. The conditions that maximize the Network Index Score no longer align with those that maximize the product of memory and information capacity.
